# Reduced local mechanical stimuli in spaceflight diminishes osteocyte lacunar morphometry and spatial heterogeneity in mouse cortical bone

**DOI:** 10.1101/2022.01.04.474962

**Authors:** Jennifer C. Coulombe, Zachary A. Mullen, Ashton M. Weins, Liam E. Fisher, Maureen E. Lynch, Louis S. Stodieck, Virginia. L Ferguson

## Abstract

Three-dimensional (3D) imaging of osteocyte lacunae has recently substantiated the connection between lacunar shape and size, and osteocyte age, viability, and mechanotransduction. Yet it remains unclear why individual osteocytes reshape their lacunae and how networks of osteocytes change in response to local alterations in mechanical loads. We evaluated the effects of local mechanical stimuli on osteocyte lacunar morphometrics in tibial cortical bone from young female mice flown on the Space Shuttle for ∼13 days. We optimized scan parameters, using a laboratory-based submicrometer-resolution X-Ray Microscope, to achieve large ∼ 0.3 mm^3^ fields of view with sufficient resolution (≥ 0.3 μm) to visualize and measure thousands of lacunae per scan. Our novel approach avoids large measurement errors that are inherent in 2D and enables a facile 3D solution as compared to the lower resolution from benchtop micro-computed tomography (CT) systems or the cost and inaccessibility of synchrotron-based CT. Osteocyte lacunae were altered following microgravity exposure in a region-specific manner: more elongated (+7.0% Stretch) in predominately tensile-loaded bone as compared to those in compressively-loaded regions. In compressively-loaded bone, lacunae formed in microgravity were significantly larger (+6.9% Volume) than in the same region formed on Earth. We also evaluated lacunar heterogeneity (i.e., spatial autocorrelation of lacunar morphometric parameters) via kriging models. These statistical models demonstrated that heterogeneity varied with underlying spatial contributors, i.e. the local mechanical and biological environment. Yet in the absence of gravitational loading, osteocyte lacunae in newly formed bone were larger and were collectively more homogenous than in bone formed on Earth. Overall, this study shows that osteocyte reshape their lacunae in response to changes, or absence, in local mechanical stimuli and different biological environments. Additionally, spatial relationships among osteocytes are complex and necessitate evaluation in carefully selected regions of interest and of large cell populations.

**GRAPHICAL ABSTRACT:** 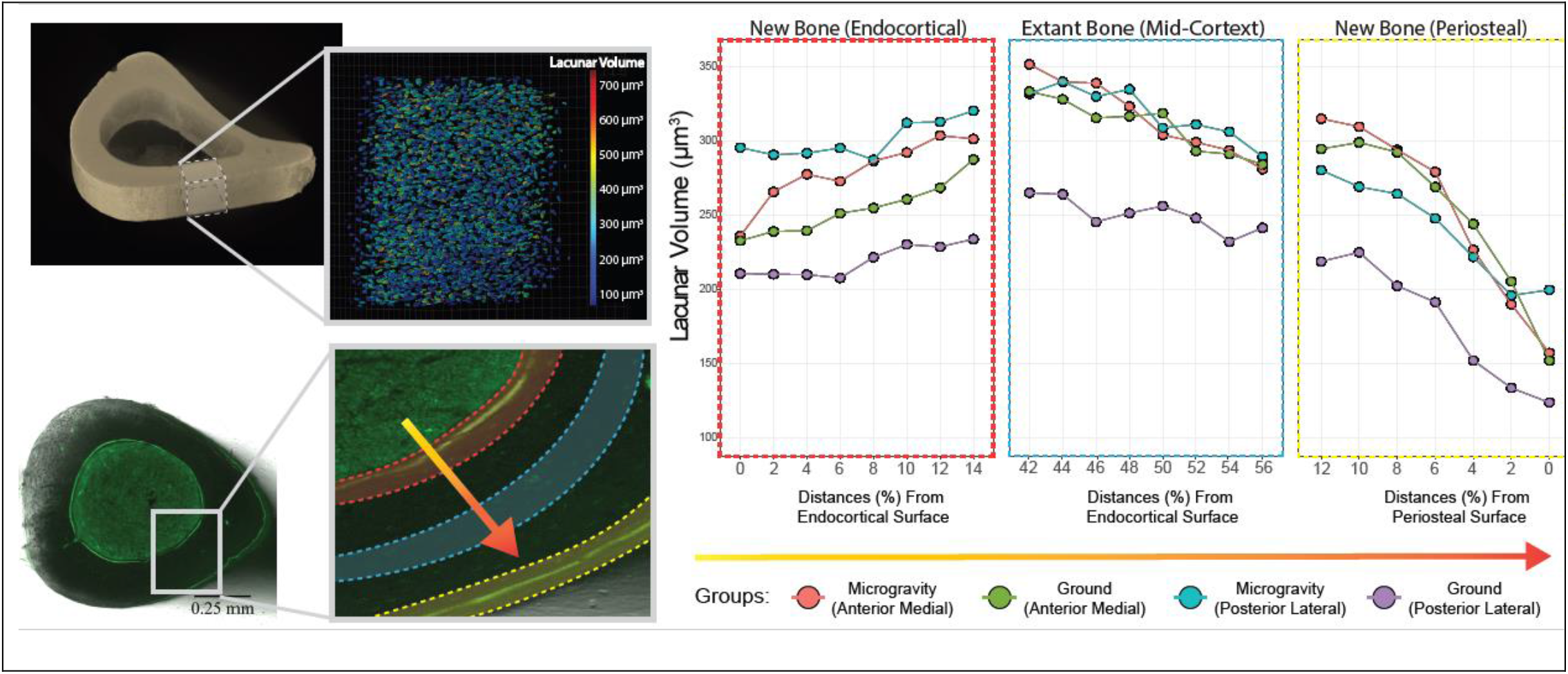

## 1. INTRODUCTION

Bone is an endocrine organ where osteocytes, the most abundant bone cells^(1)^, play a central role in communicating with other cells and tissues throughout the body. Previously, osteocytes were thought to be inert as they are embedded in hollow spaces, or lacunae, throughout the bone tissue. In the past two decades, their crucial role in mineral homeostasis, strain sensing, and paracrine signaling has been elucidated^(1)^. The complex osteocyte-lacuna-canalicular network (LCN) allows these cells to carry out their sensing and signaling role, and is crucial for maintaining mineral homeostasis and overall skeletal health. Osteocytes can signal to other cell types (i.e., osteoblasts, and osteoclasts) to (re)model bone^(2,3)^, and also change the local bone composition through alterations of extracellular matrix and mineral organization and chemistry^(4–8)^. During extreme shifts in mineral homeostasis, the volume of the osteocyte lacuna increases, such as lactation^(4,8–12)^ and hibernation^(13,14)^, via a process called perilacunar remodeling (PLR). Imbalance from pathologies, such as chronic kidney disease^(15)^, osteoporosis^(16)^, and aging^(15,17,18)^, detrimentally affect the osteocyte lacunar mineral density and morphometry to result in smaller and more spherical lacunae. In contrast, pathologies such as rickets^(15,19,20)^ osteomalacia^(19)^, osteopenia^(16)^ and vitamin D deficiencies^(21)^ along with fracture^(22)^ have larger, less uniformly oriented osteocyte lacunae. Consequentially, lacunar shape and size are sensitive biomarkers of bone’s metabolic state and systemic mineral homeostasis.

The link between the cell’s mechanosensory function, perilacunar remodeling, and lacunar morphology remains underexplored. Three key questions underpin our investigation of osteocyte function and lacunar morphology: 1) how does lacunar shape influence mechanosensation, 2) how does native mechanical loading affect lacunar shape, and 3) how do altered mechanical stimuli change lacunar shape. However, due to bone’s composition (i.e., collagen fibers and densely packed hydroxyapatite crystals), imaging and evaluating osteocyte function is challenging. Without a technique to assess how osteocytes change their lacunae *in vivo*, researchers have investigated osteocyte mechanosensitivity using *in vitro* and *ex vivo* approaches. *Hemmatian et al*., demonstrated osteocytes in larger lacunae have a stronger response to mechanical loading than osteocytes in smaller, similarly shaped lacunae^(9)^. Theoretical and finite element models have demonstrated that changes in lacunar volume and shape alter the strain and fluid flow stimulation of the osteocyte cell body^(9,11,23,24)^. Notably, *McCredie et al*., modeled longer, thinner osteocyte lacunae that generated higher maximum principal strains on the cells than a rounder lacuna model. However these studies focused on individual osteocyte PLR in response to mechanical cues rather than changes to highly interconnected populations of these cells. Future studies that leverage thousands of osteocyte lacunar morphometrics may reveals new insight into how osteocytes collectively respond local mechanical stimuli and biological environmental factors.

While mechanical stimulation is critical for osteocyte viability^(25)^, whether osteocytes reciprocally adapt their lacunae in response to novel stimuli is poorly understood. During development, the presence or absence of mechanical loading has been shown to direct osteocyte shape; osteocytes in 6-week-old mice subjected to mechanical loading were flatter and spindle-shaped, whereas osteocytes in neurectomized mice were round and irregularly shaped^(26)^. While osteocytes are mechanosensitive, simply unloading bone does not generate a uniform response. Mice exposed to 30 days of microgravity had smaller, more spherical lacunae in the femur^(27)^, yet a separate study demonstrated after 15 days of microgravity osteocyte lacunae in the mouse iliac crest were more circular and had greater area than controls^(28)^. Moreover, Earth-based disuse models have not demonstrated detectable differences in lacunar morphologies^(29)^. With new spaceflight missions and an increasingly sedentary population, determining if osteocyte lacunar morphologies are an adaptive response to disuse may be key for designing future therapeutics. Yet why, and how these morphometric differences are mediated *in vivo* is not well understood as well as whether osteocyte shape and size change with novel mechanical stimuli.

Each lacuna possesses a complex three-dimensional shape that is not adequately or consistently captured in 2D. Osteocyte lacunae have been largely evaluated for morphometry using two-dimensional measures, including histological stains, TEM, and SEM^(6)^. Though these techniques provide high-resolution images, there are inherent biases in 2D lacunar measures compared to 3D. For example, 2D measures of lacunar area, perimeter, circularity, and density are poor predictors of 3D measures of lacunar volume, surface area, sphericity, and 3D density, respectively^(17)^. With advances in technology, a variety of 3D methodologies have arisen. Briefly, 3D methods including confocal laser scanning microscopy (CLSM) and multiphoton microscopy can visualize the osteocyte, the dendrites, the lacunae, and the canaliculi^(18,30)^. These microscopy techniques are limited to a small field of view (FOV) of the LCN at shallow tissue depths. By contrast, x-ray-based synchrotron radiation computed tomography (SR-CT) can capture thousands of osteocyte lacunae with resolutions as high as 30 nm in larger volumes (< 100 µm^3^ FOV) of bone than CLSM^(31–33)^. However, SR-CT is not widely accessible: beamline facilities are expensive to build and maintain, and preparations to use a beamline can take over a year^(33,34)^. Recently, commercially available x-ray based ultra-high-resolution desktop micro-computed tomography (µCT) and X-ray microscopes (XRM) have become more accessible, with comparable resolution range as SR-CT, and the potential to elucidate cellular structures like CLSM^(35)^. However, even within the same instrument, there are a variety of parameters and measures that are reported when assessing osteocyte lacunar morphometries^(17,24,36,37)^. While these technologies improve our ability to assess lacunar morphometries, they also bring a unique challenge: XRM, µCT, and SR-CT are capable of imaging up to hundreds of thousands of osteocyte lacunae in a single scan and produce massive datasets. To appropriately manage large heterogeneous datasets, conventional statistical analyses are limited and, instead, modern statistical approaches are necessary that leverage spatial correlations are necessary. Through the characterization of the morphometries of osteocyte lacunae populations, we can investigate how osteocyte networks respond to novel stimuli *in vivo*.

In this study, we demonstrate an approach for evaluating thousands of osteocytes within a large region (∼ 0.3 mm^3^ FOV) and resolution sufficient to accurately visualize and measure lacunae (≥ 0.6 μm) in mouse cortical bone tissues using a laboratory-based X-Ray Microscope. We apply these optimized parameters to evaluate how local variations in mechanical stimuli influence mouse cortical bone osteocytes to alter their lacunar morphometry when subjected to musculoskeletal unloading in spaceflight. Additionally, we demonstrate an underlying spatial signal to morphometric measures in populations of osteocyte lacunae that may explain one source of lacunar morphometric heterogeneity.

## 2. METHODS

### 2.1 Specimen

#### 2.1.1 X-Ray Microscope Parameter Optimization

Tibiae of a 12-month-old female BALB/C mouse from a cohort for an aging study were harvested and cleaned of all non-osseous tissue. The tibiae were wrapped in PBS-soaked gauze and kept at -4 degrees. All procedures were approved by the University of Colorado Animal Care and Use Committee (IACUC).

#### 2.1.2. Effect of Short-Term Microgravity Exposure on Lacunar Morphometry

Female 9-week-old C57BL/6N mice from Charles River Laboratories were launched from Kennedy Space Center on August 8, 2007, as part of the Space Shuttle mission STS-118 (STS-118). Matched ground-controls (GC) were housed in an environmental chamber mimicking the Shuttle mid-deck environment. All mice were acclimated to NASA Animal Enclosure Modules (groups of n = 4/partition) with food in the form of NASA nutrient-upgraded rodent food bars (NuRFB, Harlan Teklad TD 04197) and water Lixits® provided ad libitum for 2 weeks before the start of the study. All mice were maintained on a 12:12 hr light:dark cycle and administered a single injection of a calcein green (20 mg/kg, s.c.) one day prior to launch; mice were flown onboard the Space Shuttle Endeavour and the International Space Station for 12.8 days. Upon return, Microgravity mice were euthanized within 3-6 hours of landing via sedation with CO_2_, followed by exsanguination and cervical dislocation. Ground control mice were subjected to the same protocols as the Microgravity group on a 2-day delay due to timing constraints surrounding shuttle launch and return. All groups were sized n=12 mice/group; mice were randomly assigned to Ground Control or Microgravity groups. Tibiae were fixed in 10% neutral buffered formalin for 2 days and preserved in 70% ethanol before imaging. All procedures were approved by the Institutional Animal Care and Use Committees at the University of Colorado-Boulder; protocols for the NASA STS-118 mission were also approved by the ACUC at NASA Ames Research Center and NASA Kennedy Space Center.

### 2.2 Osteocyte lacunae visualization and quantification

The posterior lateral aspect of the proximal tibia of a 12-month female BALB/C mouse was scanned 5 mm from the tibiofibular junction (TFJ) using an Xradia Versa 520 X-Ray Microscope (‘XRM’, Carl Zeiss, Germany) to create computed tomography at sub-micrometer length scales. From this region of the bone, for an approximate volume of 0.3 mm^3^, we segmented osteocyte lacunae and quantified their morphometries. We evaluated the effects of XRM resolution, power, and voltage, and source to detector distance on the visualization and quantification of osteocyte lacunae to determine ideal imagining parameters. We sought to identify parameters that minimized scan time while producing scans with a greater signal-to-noise ratio for reliable segmentation and morphometric analysis of osteocyte lacunae. Then we described osteocyte lacunar morphometries in mice using the parameters previously identified to quantify the effects of microgravity exposure and anatomical location on osteocyte lacunar morphometry.

Samples were mounted in custom sample holders and initially visualized with the 0.4ⅹ objective to view the entire tibia. Using the TFJ as a landmark, regions of interest were identified (**Fig. 1.A**). A full reconstruction of the region of interest was created using the 4ⅹ objective that captured a volume of 3.9 mm^3^ (**Fig. 1.B**). From the 4ⅹ reconstruction, a subsection was selected based on anatomical orientation (i.e. posterior lateral or anterior medial) and scanned using the 20x objective capturing a 0.3 mm^3^ volume of bone (**Fig. 1.C**). These scans were then segmented, and osteocyte lacunae were visualized from within the bone (**Fig. 1.D**).

**Figure 1.**
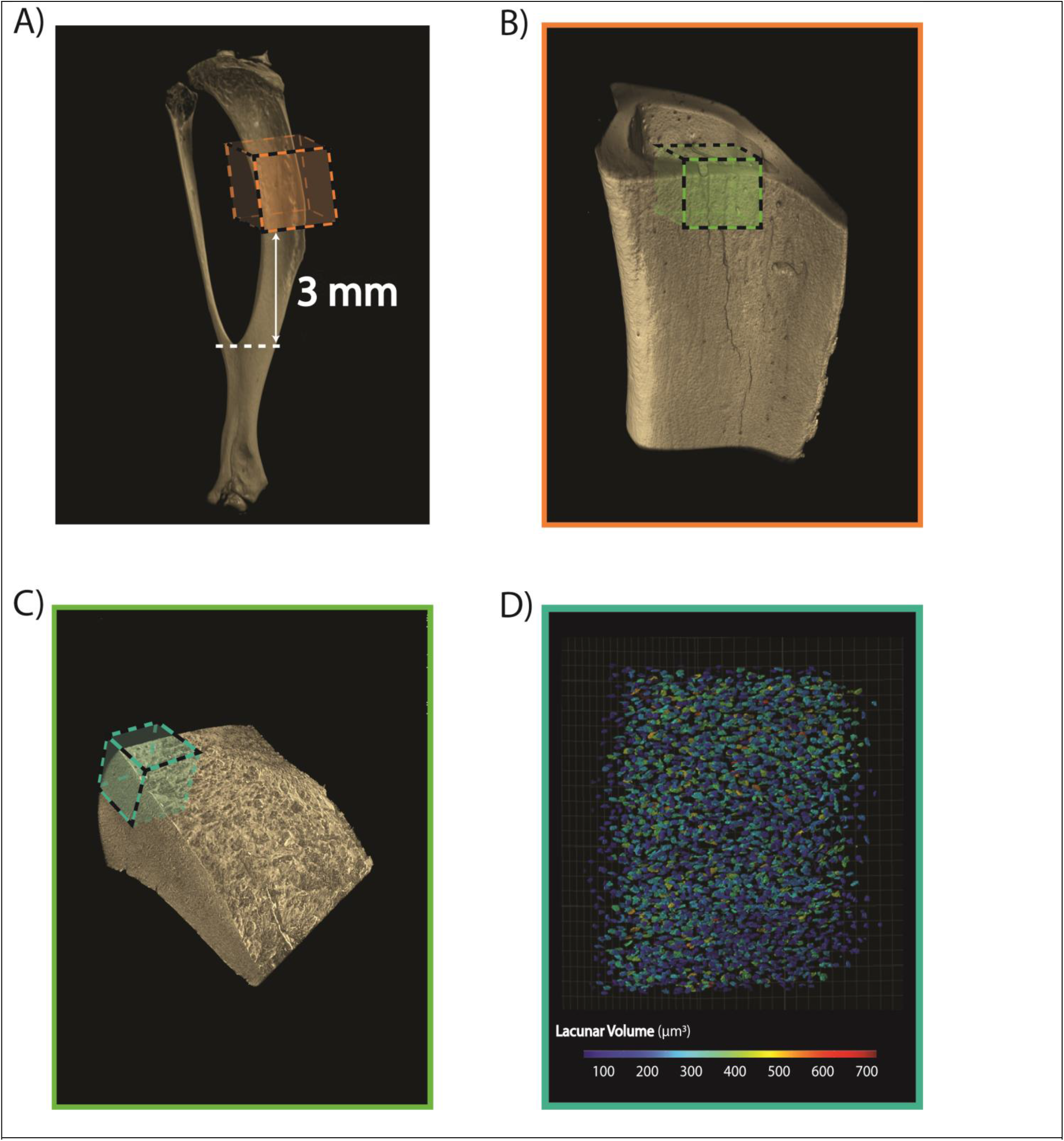
3D visualization of a mouse tibia at different objectives using an X-Ray Microscope. **A)** Representative reconstructed 0.4x objective scan of the entire tibia. **B)** Reconstructed 4x objective scan of bone region 3 mm from the tibiofibular junction. **C)** Reconstructed 20x objective scan of a 0.3 mm^3^ field of view of the bone region in the anterior medial aspect of the tibia. **D)** Negative image provides a visualization of osteocyte lacunae in bone, where lacunae are false colored by lacunar volume. Scans were reconstructed and visualized using Dragonfly 2020.2 Pro [Object Research Systems (ORS) Inc, Montreal, QC].

To determine the number of projections for each scan we used the following equation:

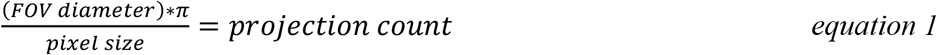

Where FOV represents the field of view and assumes isometric voxels. This equation provides a projection count such that there are enough voxels to outline the outer diameter and capture the entire sample. This provides enough projections to capture the entire sample, and might help improve the signal-to-noise ratios.

### 2.3 Evaluation of Geometric Magnification and Nominal Resolution on Lacunar Morphometry Precision

The appropriate resolution for nano-computed tomography (nanoCT) of osteocyte lacunar morphometric characteristics using an XRM was determined through repeated scans of the same region of bone. The posterior lateral aspect of a proximal tibia was scanned at 3 mm proximal to the TFJ with six different resolutions: 5 µm, 2.5 µm, and 1.2 µm using a 4ⅹ objective, and 1.2 µm, 0.6 µm and 0.3 µm using a 20x objective. The source and detector distance were varied between 16.5 and 19.3 mm from the sample, and exposure ranged from 1 - 40 seconds contingent on resulting intensity values. 3201 projections were collected for each of the scans. Scans from the 4ⅹ objective were sub-sectioned to include only the region of interest captured by the 20x objective scans for a more equivalent comparison of lacunar measures. Outcomes are reported in **Supplemental Document Table 1, Supplemental Figure 1, and *§IV. II***.

### 2.4 Evaluation of Power and Voltage on Lacunar Morphometry Precision

The effect of power and voltage on osteocyte lacunar measures was assessed in the anterior medial aspect of a proximal tibia from a 12-month-old female BALB/C mouse was scanned 3mm from the TFJ with matched parameters: 20x objective, 0.6 µm resolution, with 3201 projections, 13.1mm between source and detector. The power and voltage varied between scans, 40kv, 3W; 60kv, 5W, 80kv, 7W; 100kv, 9W and 140kv, 10W with exposure ranged from 0.7 to 9.5 seconds per projection contingent on resulting intensity values. Outcomes are reported in **Supplemental Document Table 2, Supplemental Figure 2, and *§IV. III***.

### 2.5 Evaluation of Source and Detector Position on Lacunar Morphometry Precision

Two scans of the same region of interest in the same bone were taken with matched parameters: 40kv, 3W, 20x objective, 0.6 µm resolution, with 3201 projections, but with varied source and detector positions. The first scan had 13.1 mm between the source and detector, with an exposure time of 9.5 seconds per projection for 3201 projections. The source to detector distance was then increased to 23.1 mm with an exposure of 31 seconds contingent on resulting intensity values. Outcomes are reported in **Supplemental Document Table 3, and *§IV. IV***.

### 2.6 Evaluation of Segmentation and Quantification on Lacunar Morphometry Precision

Individual scans were reconstructed using Zeiss Scout-and-Scan Control System Reconstructor 14.0.14829.3814 (Zeiss, Dublin, CA, United States of America) then imported into Dragonfly 2020.2 Pro [Object Research Systems (ORS) Inc, Montreal, QC] in which files were segmented using a global Otsu thresholding algorithm. The segmentation yields two regions of interest (ROI): one of the bone tissues and the other of the void spaces within the bone. The void space ROI was then converted into MultiROIs using 26-connectivity that subdivided the original ROI into distinct ROIs for each void. A volume filter is used to remove any ROI less than 50 µm^3^ and greater than 1500 µm^3^, as per literature values of osteocyte lacunae^(17,24,32,36)^. Surface Area, Volume/Surface Area, Aspect Ratio, and Centers of mass in the X, Y, and Z coordinates were assessed in Dragonfly for the remaining voids following the volume filter. We developed a custom Python plugin **(Sup. *§V*)** to evaluate lacunar morphometries based on the work from *Mader et al*.^(36)^, *Heveran et al*.^(17)^, and *McCreadie et al*.^(24)^. To summarize, an inertia matrix is computed based on the center of mass for each ROI. Then the eigenvalues and eigenvectors of the matrix are calculated to determine the principal moments of inertia and principal axes directions. Assuming an ellipsoidal shape for each lacuna, the principal moments of inertia are related to the ellipsoidal radii **(Fig. 2A)**. From these ellipsoidal radii Lacunar Length, Lacunar Height, Lacunar Width and subsequently Lacunar Stretch (Lc. St., **Fig. 2C, F, G**), Lacunar Oblateness (Lc. Ob., **Fig. 2B, D, E**) are calculated. Additionally, we introduced novel measures of Fitted Ellipsoid Volume, Fitted Ellipsoid Surface Area, Lacunar coefficient of variance, and Signal to Noise (SNR). Through comparisons of Lacunar Volume and Surface Area, these measures provide an approximate amount of error for the lacunar measures that assume an ellipsoidal shape. SNR was calculated by dividing the mean intensity value (in arbitrary XRM units of intensity) of the entire volume of the bone by the mean intensity value of the lacunae. Similarly, the lacunar coefficient of variance was calculated by dividing the mean intensity of each lacuna by its respective standard deviation of intensity. These two measures provide values to assess different scan parameters for their reliability of segmentation; a higher SNR indicates a better contrast between the lacunae and the bone, and a lower lacunar coefficient of variance value indicates less noise in each lacuna. Bone volume (BV), Tissue volume (TV), Number of Lacunae (Lc. N), and Lacunar Density (Lc. D), and duration (in hours) were also evaluated for each scan.

**Figure 2.**
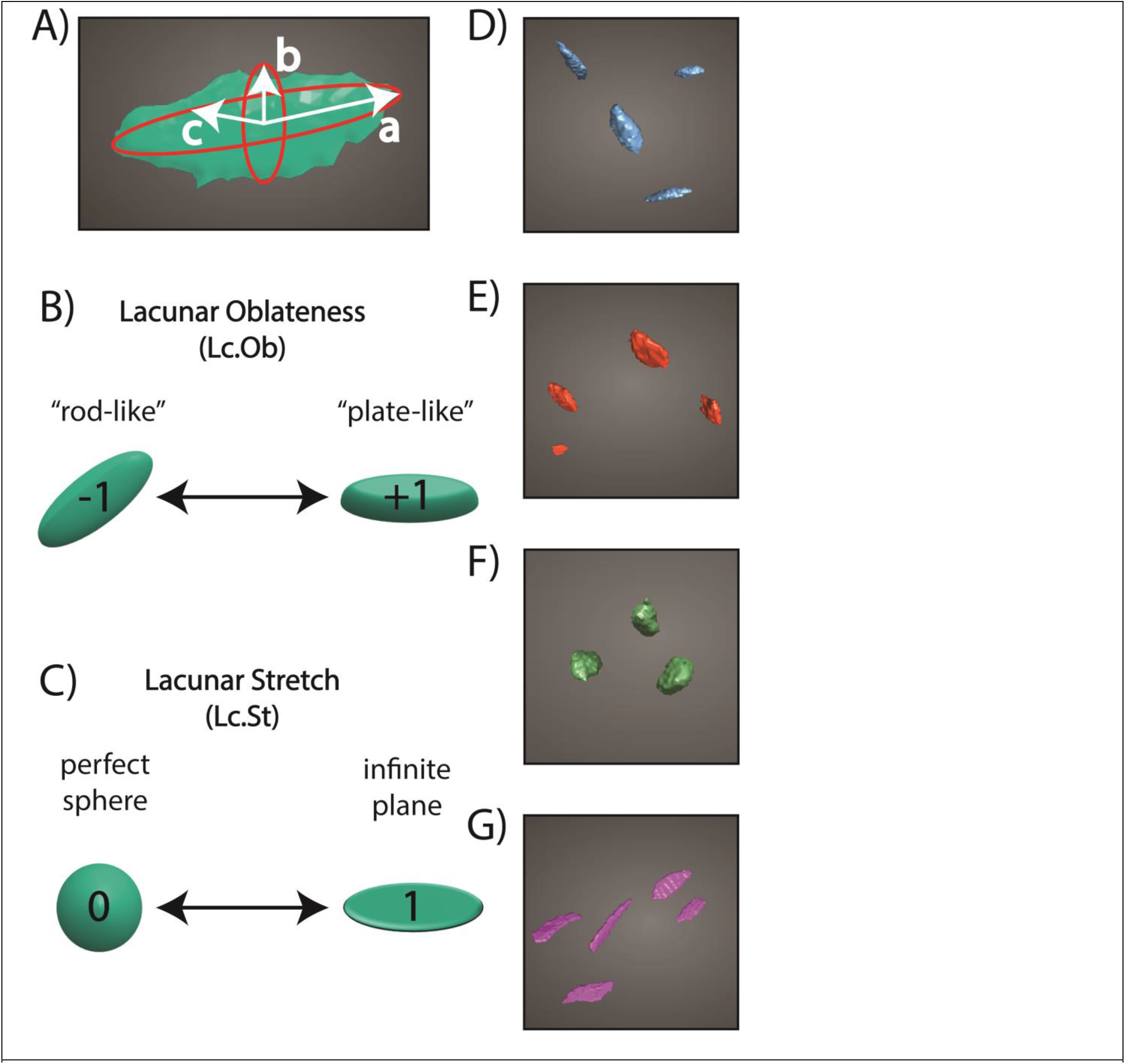
Osteocyte lacunar shape parameters. **A)** Representative osteocyte lacunae fitted with ellipse (red) with the ellipsoidal radii represented with a white arrow and defined as (a) Lacunar Length, (b) Lacunar Height, and (c) Lacunar Width. Shape measures of the lacunae adapted from *Mader et al*.^(36)^, including **B)** Lacunar Oblateness (Lc. Ob), a range of values to describe the “plate-ness” of an ellipsoidal shape and **C)** Lacunar Stretch (Lc. St.), a range of values to describe the sphericity of a shape. Representative osteocyte lacunae with comparable volumes (∼200 µm^3^) illustrating range of oblateness values **D) & E)** and stretch values **F) & G). D)** “Rod-like” lacunae with oblateness values of -0.95. **E)** “plate-like” lacunae with oblateness values of +0.95. **F)** Spherical lacunae with stretch values of +0.35. **G)** Planar lacunae with stretch values of +0.9.

### 2.7 Effect of Short-Term Microgravity Exposure on Lacunar Morphometry

Following our assessment of optimized imaging parameters for a medium through-put presented in §2.2-2.6, we applied our method for visualization and quantification of osteocyte lacunar morphometries in different microgravity exposure conditions and anatomical location in young, growing female mice.

Samples were scanned, while in 70% EtOH, in custom-made sample holders that oriented the samples vertically on the stage. Tibiae were originally scanned with a 4ⅹ objective to determine regions of interest. A 0.3 mm^3^ region of bone was located on the anterior medial and posterior lateral aspect of the tibia, 3 mm from the TFJ. These locations were selected from FE models of in vivo strain from previous work, where anterior medial and posterior lateral experienced primarily tensile or compressive strain, respectively^(38)^. Nano-computed tomography images image were collected in the region of interest using a 20x objective with energy settings of 40 V, 3.0 W, using the Air Filter and 3,201 projections. To obtain a constant resolution of 0.6 µm voxel for all the samples, the source and detector distance were varied between 5.8 and 8.5 mm from the sample, and exposure ranges from 6 - 8 seconds contingent on resulting intensity values.

Regions of interest were analyzed in Dragonfly 2020.2 Pro. Lacunae were segmented and quantified as previously described presented in §2.6. Additionally, the minimum and average distance between each lacuna and the periosteal and endocortical bone were calculated. Using a Sobel filter, an image processing edge detection algorithm, in Dragonfly Pro 2020.2 the periosteal and endocortical surface of the bone were automatically identified and segmented. For each osteocyte lacuna, the minimum distance from the endocortical and periosteal surface was calculated. For each sample (i.e. Flight Tibia 1 Anterior Medial, Flight Mouse 1 Posterior Lateral, Ground Tibia 1 Posterior Lateral, etc.) distances from lacunae to bone surface were normalized to the sample width. Osteocyte lacunae were then binned by their distance (in percent) from both the endocortical and periosteal surface.

“New” bone extended from either the endocortical or periosteal bone edge to fluorescent calcein labels (**Fig. 3A**). Extant bone was conservatively defined as the mid-cortex or the 50% distance from either the endocortical or periosteal surface. 8 bins, corresponding to 16% of the total width of the bone, were compared for each region (i.e. new bone at endocortical surface, extant bone, new bone at the periosteal surface), capturing a representative 48% of the total volume of the bone (**Fig. 3B**).

**Figure 3.**
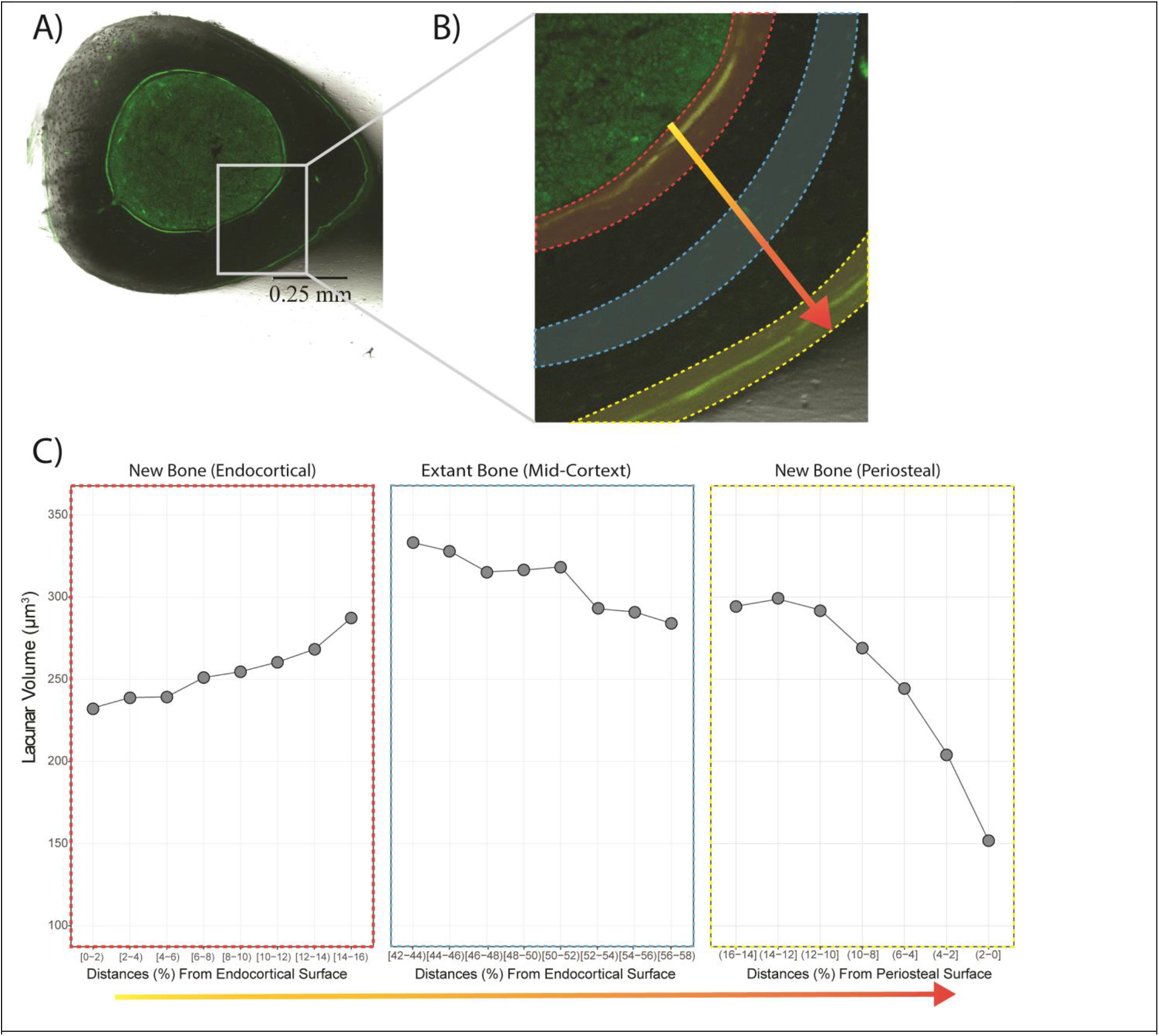
Osteocyte lacunae binned by distance from endocortical and periosteal surfaces. **A)** Representative Brightfield image of a cross-section of the left tibia with fluorescent calcein labels for quantitative histomorphometric measures of bone metabolism. **B)** Blown-up image of the anterior medial region of the bone. False-colored regions mark (red) new bone at the endocortical surface, (blue) extant bone around 50% of the total width of the cortical region, and (yellow) new bone at the periosteal surface. Arrow denotes the direction of the bins evaluated. **C)** Representative plots of the volume of lacunae in each bin in three different regions of the bone. Binning was based on the percent distance of the total width of each bone sample. Each bin represents 2% of the total bone width.

**Figure 4.**
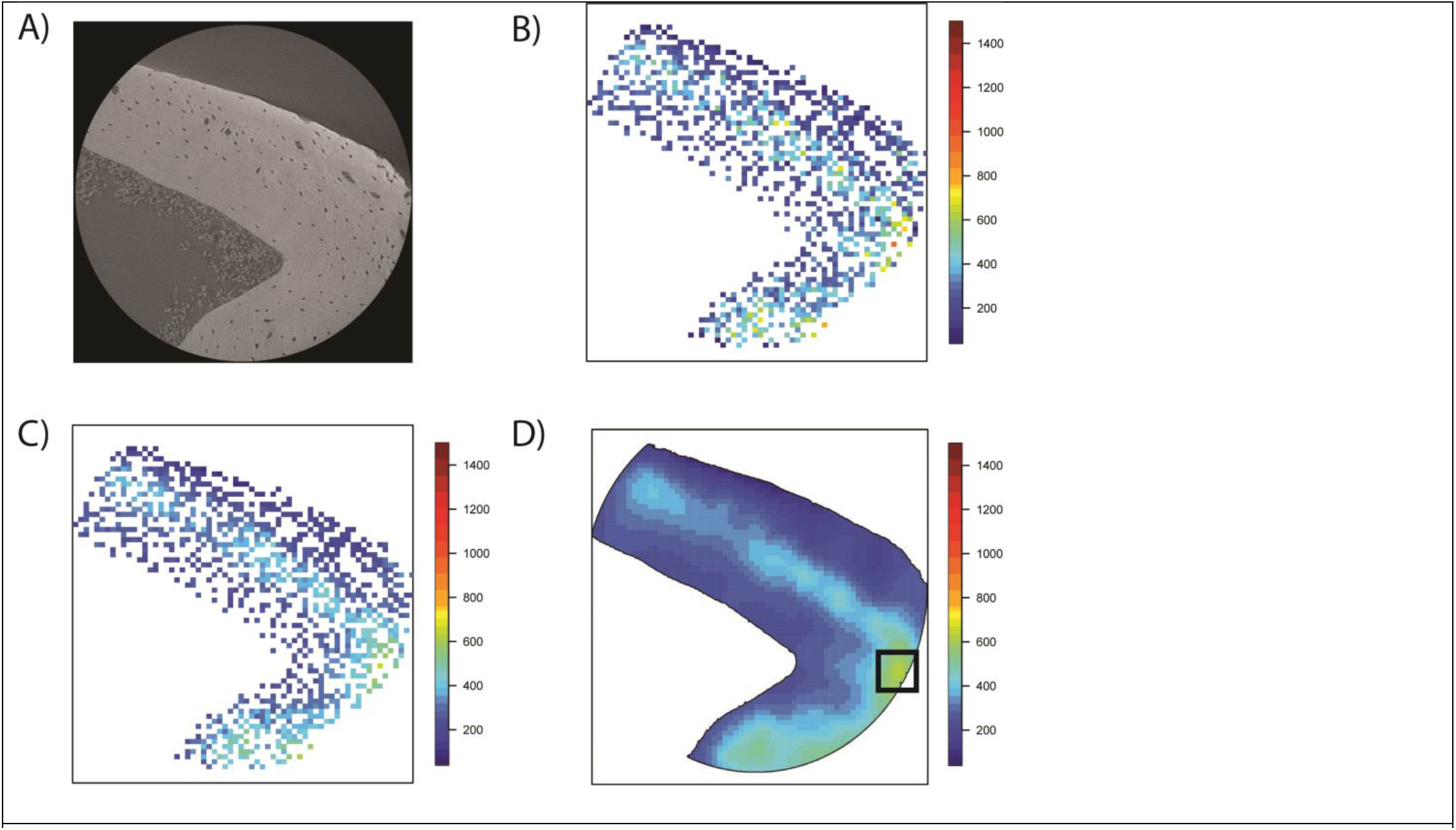
Steps of kriging model of lacunar volume construction. **A)** XRM cross-section of representative posterior lateral sample. **B)** Raw values of lacunar morphometrics represented by colored squares. X and Y axis are arbitrary coordinate grid from XRM image. **C)** Kriged values of lacunar volume projected through ∼40% of total volume of the bone centered at the midpoint in the Z-axis. The “mean” response from B, has removed the effects of Sigma from the raw data, smoothing out extrema. **D)** Final Kriged predictive model of lacunar volume for infinitesimal plane at exact center of the Z-axis. Black box encompasses a “spatial feature”, or a region of spatial autocorrelation that is visually identifiable from surrounding regions.

### 2.8 Statistics and Data Analysis

#### 2.8.1 Summary Statistics

Data sets were evaluated for normality, skew, kurtosis, and heteroskedasticity. Data are presented as the median and interquartile range (IQR) for all data sets that were skewed and failed to meet normality assumptions. The effects of resolution, power, and voltage, and geometric magnification on osteocyte lacunar morphometric measures were evaluated for each parameter using ANOVA or Mann–Whitney U test as appropriate. The Tukey posthoc procedure was utilized to compare means between levels within each parameter. Multifactor repeated measures ANOVA with Tukey’s HSD Post Hoc analysis assessed the effect of microgravity exposure, anatomical location, and bone region on osteocyte lacunar morphometric characteristics. For all analyses, alpha was set a priori to p = 0.05. All statistics were performed using R (Version 3.5.1).

#### 2.8.2 Kriging Models for Evaluating Spatial Statistics

If we consider lacunar osteocyte networks as responding to an unobserved underlying continuous spatial forces, the mean response as a function of location could be well fit by a kriging estimator **(Fig. 7)**.

A kriging model proposes a general model of the form:

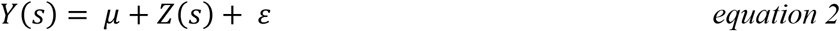

where μ is the mean osteocyte lacunar morphometric measure (i.e. stretch, oblateness, volume, surface area, etc), vectorized *Z* is a correlated Gaussian process observed at sites s_1_, s_2_, …, s_n_ and ε is a white noise process, e.g. ε ∼ *N*(0, τ^2^).

In particular, we propose ª, the isotropic exponential covariance, for its ease of computation and frequent use in spatial literature. This results in an underlying density of:

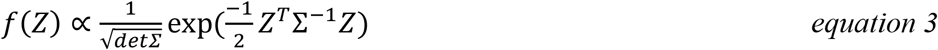

With covariance matrix Σ given by the Euclidean coordinates of the data following an exponential process:

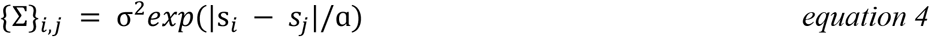

Such a covariance function captures information including the effective distance at which osteocyte lacuna exhibit 50%, 10%, etc. structural autocorrelation.

The kriging predictor smooths the observational data to predict mean morphometric measures at either existing or unobserved locations s_0_ in the bone via the estimator:

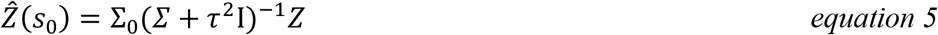

Where {Σ_0_}_i,j_ holds covariances between location *s*_0_ and data locations *s*_*i*_, as in *equation 4*. Parameter estimation of ª, τ^2^, and σ^2^ is performed by a quasi-Newton solver on the log-likelihood function (*equation 3*) in the R package optim.

ª is called the “range” parameter and controls the effective distance at which osteocytes exhibit spatial autocorrelation. τ^2^ and σ^2^ are both variance parameters that assign the variance to either unattributed noise or spatial variation.

For the non-stationarity portion of this analysis in *§3*.*3* the same optimization schema was used on a window size of 20 µm.

## 3. RESULTS

XRM imaging of osteocyte lacunae in the mouse tibia (**Fig. 1**) was enabled by optimizing the challenging trade-offs among spatial resolution, signal-to-noise ratio (SNR), morphometric measurement precision, and scan duration. We compared the effects of nominal and geometric magnification, power and voltage, and source and detector distances on lacunar morphometric measures (**Sup. *§IV***). Using optimized imaging parameters, we then compared the lacunar morphometries (**Fig. 2**) in tibia of mice exposed to 12.8 days of microgravity exposure on the STS-118 Space Shuttle Endeavor mission to matched Ground Controls to evaluate how alterations in local mechanical loading and biological environment (i.e., newly formed vs extant bone) influenced lacunar remodeling.

### 3.1 Microgravity exposure alters osteocyte lacunar morphometry

Having established histomorphometry measures from the contralateral limb (**Fig. 3A**), we sought to determine if changes in osteocyte lacunar morphologies are an adaptive response to disuse. For our lacunar analysis, we defined distance ranges from the endocortical and periosteal surfaces as either new or extant bone (**Fig. 3B & *§2*.*7***).

Osteocyte lacunae formed under microgravity conditions (i.e., in ‘new’ bone) had larger volumes, but in a site-specific manner. In the posterior lateral (PL) tibia of mice exposed to microgravity, osteocyte lacunae were larger (+6.9% Volume, p<0.001) in the newly formed bone as compared ground controls (**Fig. 5.A**). Yet, no changes in lacunar volume were observed following microgravity exposure in extant bone within the PL quadrant (**Fig. 5.A**). The same pattern was not seen in the anterior medial (AM) side of the tibia. For both Microgravity and Ground Control mice, and at both the PL and AM regions, osteocyte lacunar volume near the endocortical surface was not significantly different than lacunae at the periosteal surface.

**Figure 5.**
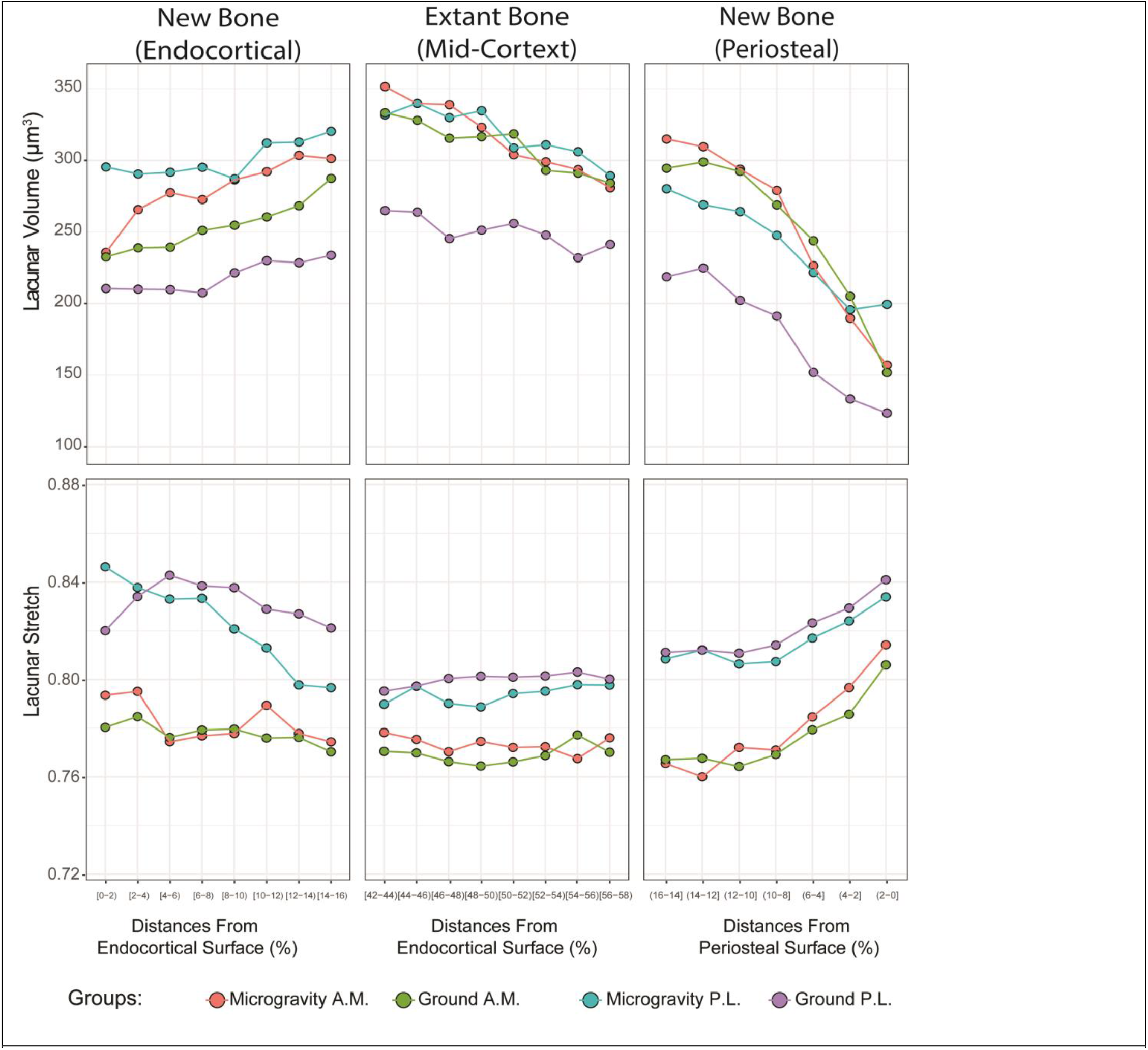
Osteocyte lacunae formed under microgravity conditions in the posterior lateral aspect of the tibia are larger than those formed at 1g. **A)** Mean lacunar volume (µm^3^) and **B)** lacunar stretch changes as a function of distance from the endocortical and periosteal bone surfaces. Distances are presented as the percent of the total width of the cortical region scanned. Data were binned in 2% ranges starting at the endocortical surface. “New” and “Extant” bone designations were determined from quantitative histomorphometry of fluorescent labels on the contralateral limb.

Unlike lacunar volume, the lacunar shape did not change with microgravity exposure. Yet the lacunar stretch was also highly site- and region-specific. Osteocyte lacunae in the PL region of both Microgravity and Ground Control tibia were more spherical (+6.7% Stretch Microgravity, p<0.001; +5.1% Stretch GC, p<0.001) than the lacunae in the AM region at the endocortical surface (**Fig. 5.B**). Additionally, lacunae in the bone near the endocortical and periosteal surfaces were more spherical than lacunae in the extant bone at PL, but not AM regions. For example, in the PL region of the Ground Control tibia, osteocyte lacunae at the endocortical surface were +3.0% more spherical (p = 0.008) than osteocyte lacunae in the extant bone at the mid-cortex (**Fig. 5.B**). A similar pattern was detected in osteocyte lacunae in Microgravity tibia. Osteocyte lacunae at the endochondral surface of the PL site were +7.0% more spherical (p < 0.001) than lacunae in the extant bone (**Fig. 5.B**). No differences in lacunar stretch were detected between lacunae at the endocortical and periosteal surfaces. Despite the site-specific differences in lacunar stretch, lacunar oblateness did not vary with either microgravity exposure or anatomical location.

### 3.2 Microgravity exposure alters lacunar spatial patterns

As lacunar morphometries were different near the surfaces of the bone as compared to the mid-cortex, we next investigated the spatial variation of volume, stretch, and oblateness within individual samples. Borrowing from geostatistics, we generated a continuous spatial model of lacunar morphometries over a sample of the bone by implementing kriging^(39)^. The kriging estimator is the best linear and unbiased estimator for a Gaussian process with a known covariance structure between observations. We explored covariance measures with three parameters of the spatial model for differences between anatomical region and microgravity exposure: range (ª), nugget (σ^2^), and process variance (τ). These covariance parameters represent bone-specific measures, such as “the proportion of variability attributable to continuous spatial variation (compared to overall variability).”

Spatial measures were determined largely by anatomical location, similar to our observations of lacunar morphometric parameters (***§3*.*1***). In the Ground Control tibiae, the range, nugget, and process variance for volume, stretch, and oblateness were significantly different in the AM region than the PL region. For example, in the PL aspect of the tibia, the shorter ranges indicated regions of autocorrelated osteocyte lacunar volumes that were smaller and more disconnected (**Fig. 6.B.)**. By contrast, in the AM aspect of the tibia, regions of higher range corresponded to bones with larger, more connected regions of similarly shaped or volumed osteocyte lacunae (**Fig. 6.B.)**. Similar to spatial models of the tibia of Ground Control mice, tibia from mice exposed to microgravity had spatial models of lacunar volume that were significantly different between the anatomical regions (**Fig. 6.B.)**, yet no differences in the spatial variance parameters of lacunar shape (stretch and oblateness) were detected.

**Figure 6.**
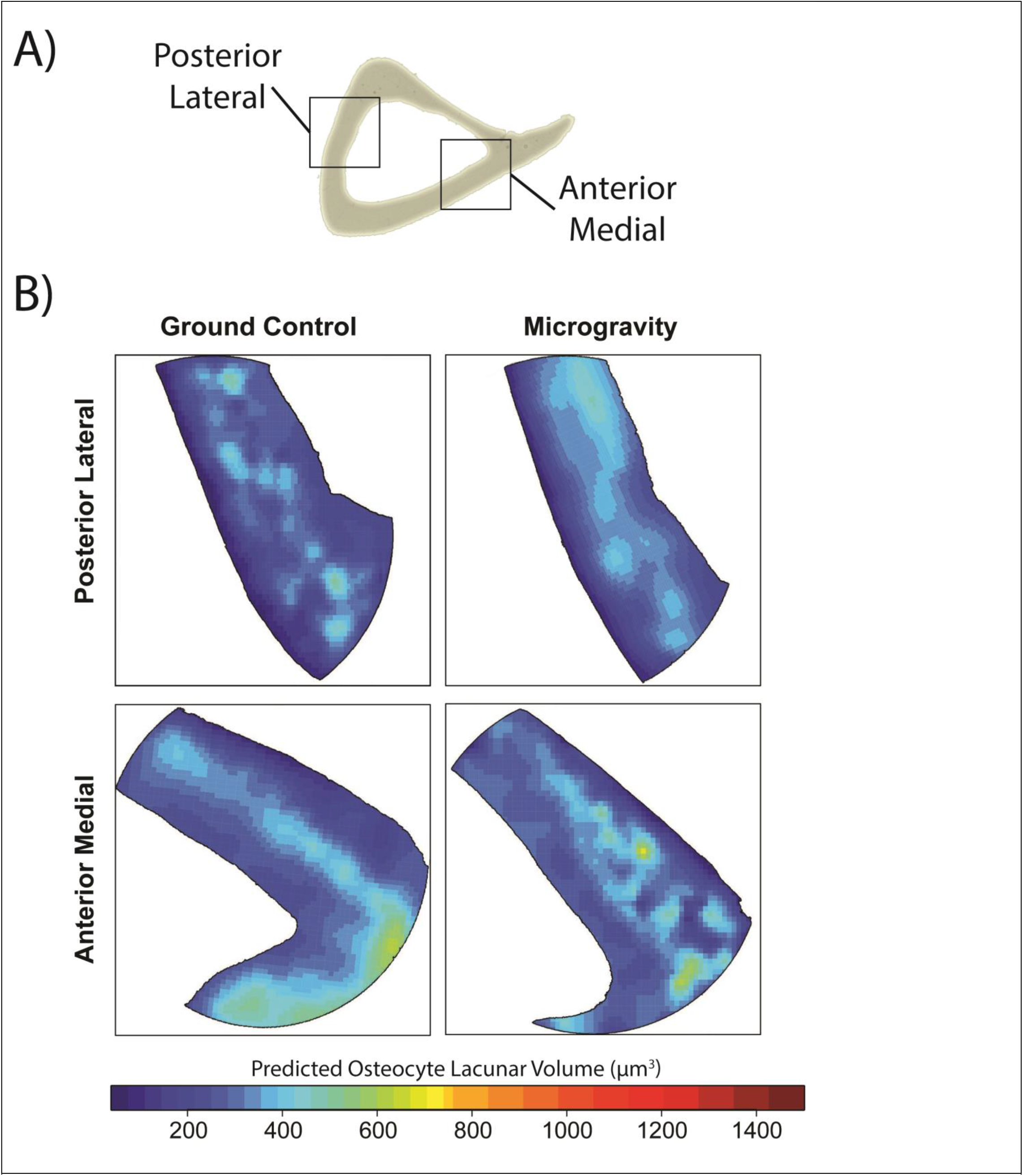
Kriging predicted mean of osteocyte lacunar volume of bisectional cut of individual bone samples. **A)** Cross-section of proximal tibia with posterior lateral and anterior medial regions of interest indicated. **B)** Representative predicted kriging of osteocyte lacunar mean volume in a cross section of the posterior lateral (top) and anterior medial (bottom) tibia of Ground Control (left) and Microgravity (right) mice.

**Figure 7.**
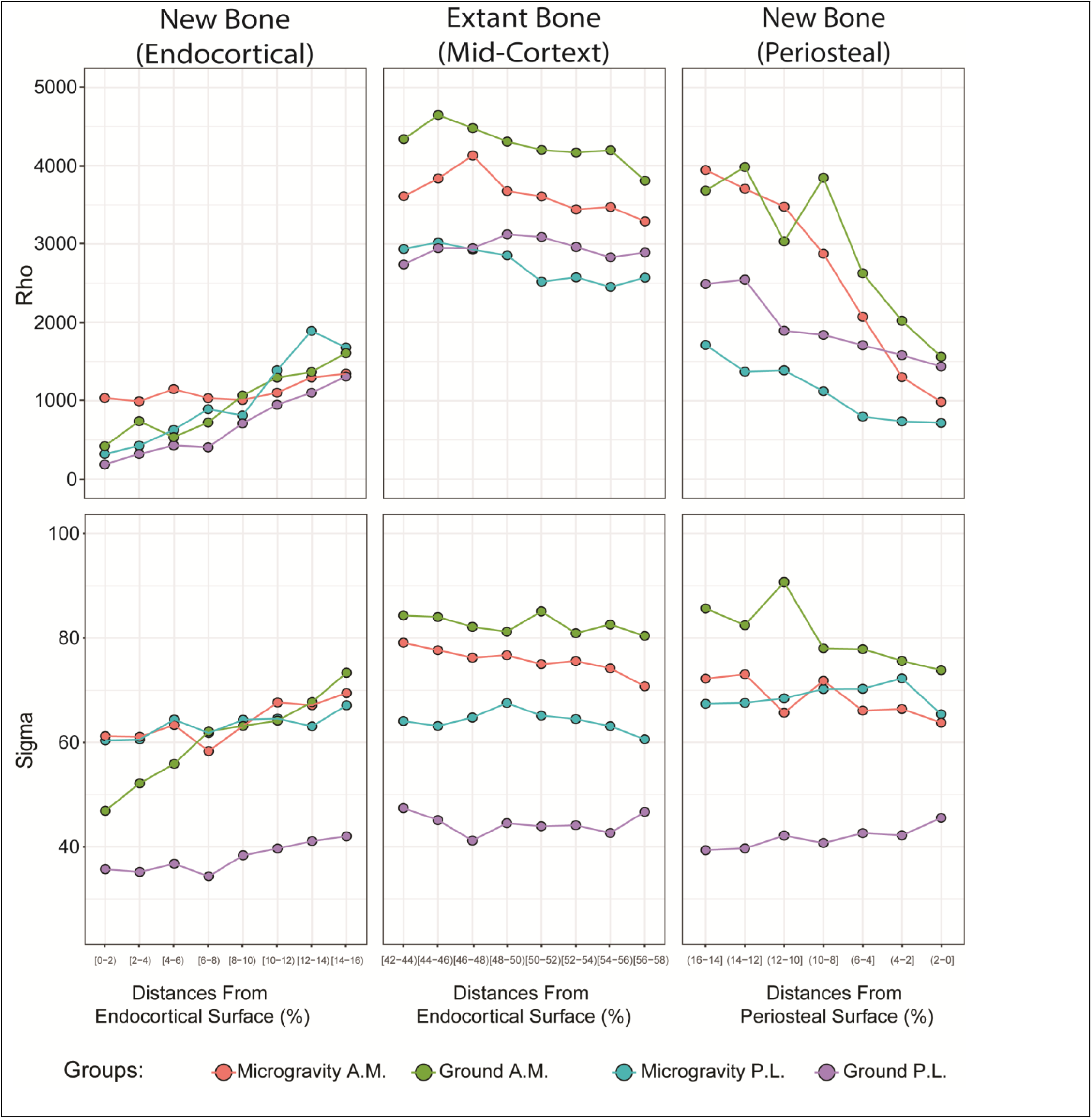
Osteocyte lacunae spatial population phenotypes are more homogenous in the posterior lateral tibia and vary by anatomical location and distance from the bone surface. **A)** Spatial parameter ρ of lacunar volume and **B)** spatial parameter σ of lacunar volume change as a function of distance from the endocortical and periosteal bone surfaces. Distances are presented as the percent of the total width of the cortical region scanned. Data were binned in 2% ranges starting at the endocortical surface. “New” and “Extant” bone designations were determined from quantitative histomorphometry of fluorescent labels on the contralateral limb.

### 3.3 Spatial lacunar populations phenotypes are site-specific

We next investigated the extent to which different regions of the bone had lacunae that had spatial autocorrelation (i.e., if osteocyte lacunae were similar to their immediate neighbors in terms of specific morphometrics). As each bone region has unique characteristics (e.g., mechanical microenvironment, composition, collagen orientation), we tested if regions may have altered “spatial population phenotypes” at different distances for the surface of the bone, and further, microgravity exposure may differentially alter those phenotypes.

The spatial model parameters described above [range (a), nugget (σ^2^), and process variance (τ)] well describe spatial features and at what distances they occur in the cross-section of the bone; however, these measures only represented a single instance of each parameter per sample. To investigate spatial autocorrelation, we allowed the spatial parameters to vary over regions within a same sample, a process known as ‘non-stationarity’. We evaluated ρ and σ of volume, stretch, and oblateness for each lacunae using a local “window” and compared these values to distances from the bone surface of that lacunae. Together, ρ and σ described the degree of spatial autocorrelation by decomposing it into two components - ρ measures the amount of variability in lacunar morphometry that is attributable to location, and σ holds the portion of the variance attributed to white noise or separate from distance.

Large ρ values indicate a perfectly smooth and continuous surface, where each individual osteocyte lacunae would be very similar to its nearest neighbors. σ describes the amount of variability we might expect between two lacunae that are arbitrarily close together. Thus, ρ and σ are negatively correlated. Performing this analysis for every lacuna provided a description for which regions of the bone exhibit more or less variation, and whether that variation was spatially correlated or not (**Fig. 4 & *§2*.*8*.*2***).

We evaluated ρ and σ of lacunar volume, stretch, and oblateness at the endocortical surface, periosteal surface, and mid-cortex. Our results suggested that there was a greater degree of spatial homogeneity in lacunar volume in new as compared to older bone. For example, at the endocortical surface, ρ of lacunar volume was less than at the mid-cortex, or the periosteal surface for all groups (**Fig. 7A**). However, at the mid-cortex, ρ of lacunar volume was greater in the AM aspect of the tibia as compared the PL aspect, regardless of microgravity exposure, indicating more homogeneity among lacunae in that region (**Fig. 7A**). A similar pattern was seen in the σ parameter of lacunar volume. Notably, σ of lacunar volume in the PL region of Ground Control mice was less than all other groups at the endocortical surface, periosteal surface, and mid-cortex (**Fig. 7B**). This result suggests that osteocyte lacunae in Ground Control mice in the PL tibia are more similar to neighboring osteocytes in terms of volume, and this spatial pattern of lacunar morphologies changes following exposure to microgravity.

## 4. DISCUSSION

Our study defined an approach for facile visualization, morphometry, and spatial assessment of thousands of osteocyte lacunae in a large FOV, which enables future studies of how individuals and networks of osteocytes undergo perilacunar remodeling. Using a laboratory-based submicrometer-resolution X-Ray Microscope, we developed a novel imaging workflow that achieved large, ∼ 0.3 mm^3^ fields of view with high resolution (≥ 0.6 μm) to visualize and measure thousands of lacunae per scan. This approach avoids large measurement errors that are inherent in 2D and enables an accessible, reliable, and facile 3D solution as compared to benchtop or synchrotron-based CT systems. We then applied our optimized imaging parameters to evaluated how individual osteocytes and populations of osteocytes remodel their lacunae in response to gravitational unloading, from 12.8 days on the Space Shuttle Endeavour, in extant and newly formed cortical bone. Osteocyte morphometric analysis and spatial statistics demonstrated that short-term microgravity exposure in young and growing female mice resulted in altered osteocyte lacunar morphometries in an anatomical site and bone region-specific manner. Overall, this work illustrated the spatial variation in osteocyte lacunar morphometries and suggests osteocyte reshape their lacunae in response to changes in local mechanical stimuli (or their absence) and different biological environments.

In this study, we found that short-term microgravity exposure in young, growing female mice resulted in lacunar morphometry changes in an anatomical region-dependent and bone site-specific manner. Osteocyte lacunae in the bone formed under microgravity conditions were larger as compared to osteocyte lacunae formed under normal 1g conditions. These differences were seen primarily in the posterior lateral side of the tibia, suggesting modulation of compressive forces may play a greater role in osteocyte lacunar morphometry than tensile strain. However, despite the volume change, no difference in shape was detected in the lacunae in the tibia of mice exposed to microgravity as seen in other studies^(37)^.

Osteocyte lacunae morphometries are highly spatially autocorrelation in a region- and microgravity-dependent manner. There was a greater degree of spatial autocorrelation (i.e., homogeneity) in lacunar morphometry measures in the posterior lateral aspect of the proximal in the ground control mice. This greater degree of homogeneity could indicate that the osteocytes in this region of the bone are under similar environmental conditions. Though subtle, this difference may suggest that in the absence of gravity and normal ambulation, osteocytes were adapting their local environment to novel (or diminished) mechanical forces. Additionally our results could suggestion under microgravity the primarily tensile or compressive regions of the proximal tibia were becoming more homogeneous in strain distributions. At a local level, site-specific differences may suggest lacunae near the endocortical, or periosteal surfaces may experience different mechanical forces or have different collagen fiber orientations than the mid-cortex of the bone. Future work will be needed to align these local spatial patterns with these underlying drivers of the spatial signal in lacunar morphometries.

The mathematical kriging estimator evaluated in this study is an answer to the question: *how does an estimate of the mean of each of our lacunar morphologies adjust to the presence of continuous spatial variationã* While previous work provided a discrete analysis of patterns in osteocyte lacunar morphometry^(36)^, our model is continuous and allows for non-stationarity, or local variations in spatial parameters (i.e. regions of higher and lower spatial autocorrelation within the same sample). Our results demonstrate that characterization of spatial patterns of osteocyte lacunar morphometries provides better insight into the heterogeneity within the population than classic parametric analysis (e.g., ANOVA, Student’s t-test). More immediately, these findings suggest that the degree of spatial autocorrelation should be considered when comparing large populations of osteocyte lacunae to improve detection of a signal from treatment from an underlying white noise process. In other words, sampling neighboring osteocyte lacunae will reduce variance in morphometry measures. However, researchers must in turn take additional care to justify and report the location of their investigations.

Site-specific variations in osteocyte lacunar morphometry suggest that perilacunar remodeling by osteocytes is highly responsive to native mechanical or material environments, but in a region-specific manner. In our study, osteocyte lacunae in the anterior medial tibial site were larger and more elongated than lacunae in the posterior lateral site of ground control mice. Previous digital image correlation maps and finite element models of mouse tibia under axial compression demonstrated compressive strain occurs at the posterior lateral region, whereas the anterior medial region experiences primarily tensile^(38,40–42)^. These site-specific differences in lacunar morphometry may be due to the predominantly compressive or tensile forces under which the osteocyte was embedded in the bone. Furthermore, it may suggest that osteocytes “tune” their immediate environment to the kind of mechanical stimulation they experience. Alternatively, osteocyte lacunae may follow from collagen fiber orientation, which also aligns in accordance with different mechanical environments^(43)^. The arrangement of collagen fibers has been linked to the orientation of osteons in humans^(44)^, thus implying a link between osteocyte lacunae orientation and the collagen matrix^(45)^. These site-specific specializations of lacunar morphometry to different collagen fibers orientations or mechanical forces may have driven the differential effect of microgravity exposure in the anterior medial and posterior lateral region of the tibia.

The site-specific nature of osteocyte lacunar morphometry differences following microgravity in our study was also observed in the femurs of mice on the Bion-M1 spaceflight mission. The posterior mid-diaphysis region of the femur of mice on the one-month Bion-M1 spaceflight mission had more than double the number of empty lacunae compared to ground controls through histological analysis and diminished lacunar volume in using SR µCT^(27)^. Smaller, more spherical osteocyte lacunae have also been observed with aging^(17)^, and may be predictive of osteocyte death and micropetrosis, the subsequent mineralization of the lacunae^(46,47)^. By contrast, an increase in lacunar volume, as seen in our study, may be emblematic of osteocyte osteolysis (reduced mineralization) as seen in lactation^(8,9,12)^, or bone metabolic disorders such as rickets^(48)^ or osteopenia^(16)^. However, the increase in lacunar volume during spaceflight may be driven by mechanosensory cues rather than mineral homeostasis. Work by *Bakker et al*., demonstrated osteocytes in larger lacunae are more mechanosensitive than those in smaller lacunae^(9)^. Additionally, while it is well known that bone mechano-adaptive response diminishes with age^(49–51)^, recent studies of short periods of disuse were able to restore the mechanoresponsiveness of aged bone^(52–54)^. It is thus conceivable that osteocytes formed under microgravity are “tuned” to weaker mechanical signals and maintain larger lacunar volume to amplify the diminished mechanical stimulation they experience during spaceflight. The difference in lacunar morphometry changes in our study and those on the Bion-M1 mission may be due to the age and sex of the mice and the duration of the spaceflight. *Gerbaix et al*., exposed 23-week-old male mice to 30 days of microgravity exposure whereas we examined the effects of 12.8 days of microgravity exposure on young, growing 9-week-old female mice. Of additional consideration, previous spaceflight studies have suggested younger mice experience both bone formation and resorption during microgravity, while in older mice bone resorption is dominant in older mice^(55)^. While our younger mice allowed us to compare osteocyte lacunae in newly formed bone, the short duration of the mission may not be enough time for osteocytes in the extant bone to change their lacunar morphologies to a detectable degree in response to disuse. Collectively, this work suggests that younger osteocytes may be more metabolically active than mature cells, and could explain some of the site-specific differences seen in our study (i.e., new periosteal and endocortical vs midcortical bone).

This study has several limitations, including small sample sizes (N = 4). Though this produced under-powered analyses for some outcome measures, the differences in lacunar morphometries between groups were still apparent. We only evaluated mouse tibiae in this study, and future work could be expanded to consider if these measures are appropriate in studies with osteons and more complex lacunar organization. While this work demonstrates the ability to visualize osteocyte lacunae, the utility of this technique may be greatly improved if visualization of osteocytes within their lacunae can be achieved and connections to cellular or molecular changes were made. However, inferences can still be made about an osteocyte’s health or function from lacunar morphology. Lastly, the resolution limits of this instrument prevent from examining changes to the canaliculi, which are crucial for maintaining mineral homeostasis and overall skeletal health.

In conclusion, our study optimized scan parameters to achieve large fields of view with sufficient resolution to visualize thousands of lacunae per scan and measured lacunar morphometric heterogeneity via spatial statistics from the tibia of mice exposed to microgravity. Overall, this study demonstrated that osteocyte reshape their lacunae in response to changes in local mechanical stimuli (or their absence) and different biological environments. Future studies implementing osteocyte imaging (i.e., histology, microscopy, µCT, etc.) should consider a selection of anatomical location, new versus extant bone, and the additional variance it may introduce into their measures. Further elucidation of the interactions of the osteocyte with its surrounding bone environment, and inclusion of lacunar morphologies in skeletal phenotyping will also require a universal standardization of these methods and terminology.

## 5. ACKNOWLEDGEMENTS

The authors would like to acknowledge that all X-ray microscopy images were acquired in the Materials Instrumentation and Multimodal Imaging Core (MIMIC) Facility, CU Boulder (RRID:SCR_019307). We are thankful for conversations with Dr. Ralph Mueller and Elliot Goff which provided conceptual input. The research was supported by the National Space and Biomedical Research Institute awards MA00002 and BL01302 (for mouse spaceflight experiments on the STS-118 mission), the National Institute of Arthritis and Musculoskeletal and Skin Diseases under the award R21AR069791 and the National Science Foundation under the award CMMI-2047187 (in support of osteocyte lacunar assessment), the Center for Advancement of Science in Space under the award GA-2016-239 (in support of bone analysis), and the National Science Foundation under the award CMMI-1726864 (funding for the Zeiss Xradia Versa X-Ray Microscope). The STS-118 experiment was also supported in part by Amgen, Inc. and BioServe Space Technologies. The content is solely the responsibility of the authors and does not necessarily represent the official views of the National Institutes of Health. The authors acknowledge support from a graduate student training grant to JCC from the NSF, IGERT 1444807, the NIH Integrative Physiology of Aging T32AG000279-16A1, and internally from the Interdisciplinary Quantitative Biology (IQ Biology) Program at the BioFrontiers Institute.

## 6. AUTHOR CONTRIBUTIONS

J.C.C., L.S.S. and V.L.F. designed the research; J.C.C. optimized imaging techniques and collected images; J.C.C., Z.K.M., A.M.W., and L.E.F. created novel code for analysis; and J.C.C., Z.K.M., and A.M.W. analyzed data. V.L.F. contributed analytic tools. J.C.C., Z.K.M., M.E.L., and V.L.F. wrote the paper. All authors reviewed and approved the paper. J.C.C., and V.L.F. take responsibility for the integrity of data analysis.

