## Supplemental Data for "Reduced local mechanical stimuli in spaceflight diminishes osteocyte lacunar morphometry and spatial heterogeneity in mouse cortical bone"

***Contents***

I. **Abbreviations**
II. **Tables**

Table 1: Lacunar Morphologies & Nominal Resolutions	Table 2: Lacunar Morphologies & Power and Voltage

Table 3: Lacunar Morphologies & Source and Detector Distance	Table 4: Lacunar Morphologies Following Microgravity Exposure

III. **Figures**

Supplemental Figure 1: Lacunar Morphologies & Nominal Resolutions	Supplemental Figure 2: Lacunar Morphologies & Power and Voltage

IV. **X-Ray Microscopy Parameter Optimization**
V. **Python Plug-In for Dragonfly**
VI. **References**

*page 2*

*page 3*

*page 4*

*page 5*

*page 6*

*page 7*

*page 8*

*pages 9-12*

*pages 14-25*

*pages 26*

I. ***Abbreviations***

Groups:

**GC** - ground control mice

**Microgravity** - mice flown on the STS-118 shuttle mission

**Young** - mice flown on STS-118, 9-weeks old at start of experiment

Anatomical Locations:

**AM** - anterior medial

**PL** - posterior lateral

**TFJ** - tibiofibular junction

Osteocyte Lacunar Morphometry Measures:

**Lc. V (µm^3^)** - Lacuna Volume

**FE Lc. V (µm^3^)** - Fitted Ellipsoid Volume

**SA (µm)** - Surface Area

**FE SA (µm)** - Fitted Ellipsoid Surface Area

**Lc. V / SA** - Volume/Surface Area

**LAR** - Lacunar Aspect Ratio

**Lc. Sph** - Lacunar Sphericity

**Lc. Ob** - Lacunar Oblateness

**Lc. St** - Lacunar Stretch

**Lc. W (µm)** - Lacunar Width

**Lc. H (µm)** - Lacunar Height

**Lc. L (µm)** - Lacunar Length

**SNR** - Signal to Noise

**Lc. N** - Lacuna Number

**Lc. N/BV (mm^3^)** - Lacunar Density

II. ***Tables***

Table 1: Comparison of osteocyte lacunar morphologies and scan durations at different nominal resolutions.

| **Resolution, Objective** | **1.2 µm, 4x** | **1.2 µm, 20x** | **0.6 µm, 20x** | **0.3 µm, 20x** |
| --- | --- | --- | --- | --- |
| **Geometric Magnification** | 2.83 | 2.26 | 2.26 | 2.26 |
| **Power and Voltage** | 40kV 3W | 40kV 3W | 40kV 3W | 40kV 3W |
| **Source and Detector Distance** | 17.2 mm | 19.3 mm | 19.3 mm | 19.3 mm |
| **Exposure, Scan Duration** | 2.0 sec, 1.8 hrs | 3.2 sec, 2.9 hrs | 10.0 sec, 8.9 hrs | 40.0 sec, 35.6 hrs |
| Lacuna Volume | 310.23 (185.44-483.54) | 111.19 (86.86-137.24) | 160.46 (131.19-186.92) | 169.86 (143.28-196.35) |
| Fitted Ellipsoid Volume | 364.41 (214.81-591.76) | 115.6 (90.78-145.21) | 177.27 (148.04-211.27) | 189.37 (161.37-225.31) |
| % Difference in Volume | -17.28 (-27.81--10.83) | -3.9 (-7.13--1.64) | -10.02 (-16.41--6.93) | -10.74 (-17.9--7.22) |
| Surface Area | 293.27 (201.03-415.5) | 140.53 (119.56-163.31) | 206.82 (179.84-235.14) | 227.89 (200.04-257.61) |
| Fitted Ellipsoid Surface Area | 281.05 (199.04-383.13) | 146.15 (124.51-170.18) | 208.91 (181.12-239.24) | 221.96 (193.71-252.98) |
| % Difference in Surface Area | 3.65 (0.23-8.11) | -3.53 (-5.52--1.86) | -0.81 (-3.05-1.14) | 2.77 (0.25-4.99) |
| Volume/Surface Area | 1.05 (0.91-1.16) | 0.79 (0.71-0.86) | 0.77 (0.7-0.83) | 0.74 (0.68-0.8) |
| Lacunar Aspect Ratio | 0.46 (0.39-0.53) | 0.26 (0.18-0.35) | 0.25 (0.18-0.34) | 0.25 (0.18-0.34) |
| Sphericity | 0.51 (0.48-0.55) | 0.54 (0.51-0.57) | 0.46 (0.43-0.49) | 0.43 (0.41-0.46) |
| Lacuna Oblateness | 0.04 (-0.21-0.32) | -0.35 (-0.58--0.07) | -0.26 (-0.51-0.03) | -0.25 (-0.5-0.04) |
| Lacuna Stretch | 0.6 (0.52-0.66) | 0.75 (0.69-0.79) | 0.79 (0.73-0.83) | 0.8 (0.74-0.84) |
| Lacuna Width | 2.66 (2.17-3.31) | 1.53 (1.33-1.78) | 1.54 (1.35-1.83) | 1.56 (1.35-1.87) |
| Lacuna Height | 4.82 (4-5.69) | 3.0 (2.6-3.46) | 3.7 (3.18-4.23) | 3.84 (3.31-4.37) |
| Lacunae Length | 6.75 (5.65-7.86) | 6.01 (5.2-6.91) | 7.32 (6.34-8.38) | 7.54 (6.55-8.66) |
| Lacunar Contrast | 63.5 | 24.0 | 14.5 | 12.8 |
| Signal to Noise | 1.10 | 1.28 | 1.27 | 1.28 |
| Lacuna Number | 3094 | 5031 | 6503 | 6617 |
| Lacunar Density | 36.3 | 62.1 | 82.0 | 88.1 |

Data are median (25% - 75% of Inter Quartile Range)

^*^*p*<0.05 compared to ground vehicle, Tukey’s HSD. α set a priori to 0.05.

*Units of Intensity are Arbitrary XRM intensity values

Table 2: Comparison of osteocyte lacunar morphologies and scan durations at different power and voltages.

| **Resolution, Objective** | 0.6 µm, 20x | 0.6 µm, 20x | 0.6 µm, 20x | 0.6 µm, 20x | 0.6 µm, 20x |
| --- | --- | --- | --- | --- | --- |
| **Geometric Magnification** | 2.26 | 2.26 | 2.26 | 2.26 | 2.26 |
| **Power and Voltage** | **40kV 3W** | **60kV 5W** | **80kV 7W** | **100kV 9W** | **140kV 10W** |
| **Source and Detector Distance** | 13.1 mm | 13.1 mm | 13.1 mm | 13.1 mm | 13.1 mm |
| **Exposure, Scan Duration** | 9.5 sec, 8.5 hrs | 2.6 sec, 2.3 hrs | 1.3 sec, 1.1 hrs | 0.8 sec, 0.7 hrs | 0.7 sec, 0.6 hrs |
| Lacuna Volume | 232.49 (179.69-314.35) | 255.77 (188-356.76) | 263.86 (182.54-373.38) | 260.69 (166.58-383.21) | 241.12 (147.94-371.3) |
| Fitted Ellipsoid Volume | 253.8 (195.69-349.84) | 288.78 (210.92-414.47) | 314.26 (215.78-453.08) | 345 (218.68-507.46) | 364.11 (220.09-576.6) |
| % Difference in Volume | -8.07 (-12.5--5.75) | -11.71 (-17.52--8.33) | -18.19 (-26.17--13.3) | -29.94 (-44.03--21.86) | -49.24 (-72.67--35.31) |
| Surface Area | 261.64 (214.55-331.64) | 304.74 (238.17-404.01) | 359.82 (265.62-483.76) | 424.19 (296.26-584.6) | 452.35 (305.17-664.61) |
| Fitted Ellipsoid Surface Area | 258.5 (210.95-326.45) | 273.32 (218.87-356.54) | 277.12 (216.19-362.64) | 286.09 (215.01-381.24) | 303.12 (219.3-414.76) |
| % Difference in Surface Area | 1.69 (-0.33-3.5) | 9.69 (5.97-14.07) | 21.52 (16.46-27) | 31.16 (25.15-36.53) | 33 (26.37-38.6) |
| Volume/Surface Area | 0.89 (0.81-0.96) | 0.83 (0.75-0.9) | 0.72 (0.64-0.8) | 0.6 (0.52-0.68) | 0.51 (0.45-0.58) |
| Lacunar Aspect Ratio | 0.2 (0.15-0.28) | 0.22 (0.16-0.3) | 0.24 (0.18-0.32) | 0.27 (0.2-0.34) | 0.27 (0.2-0.36) |
| Sphericity | 0.47 (0.45-0.50) | 0.43 (0.40-0.46) | 0.38 (0.35-0.41) | 0.32 (0.29-0.36) | 0.29 (0.25-0.32) |
| Lacuna Oblateness | -0.44 (-0.63--0.19) | -0.44 (-0.64--0.18) | -0.46 (-0.66--0.21) | -0.43 (-0.64--0.16) | -0.43 (-0.65--0.13) |
| Lacuna Stretch | 0.79 (0.75-0.82) | 0.77 (0.73-0.81) | 0.74 (0.7-0.78) | 0.72 (0.67-0.76) | 0.72 (0.67-0.77) |
| Lacuna Width | 1.86 (1.64-2.09) | 2.03 (1.75-2.29) | 2.24 (1.9-2.54) | 2.36 (1.99-2.73) | 2.37 (1.96-2.89) |
| Lacuna Height | 3.77 (3.21-4.47) | 3.87 (3.25-4.7) | 3.9 (3.26-4.72) | 4.08 (3.4-4.96) | 4.21 (3.44-5.18) |
| Lacunae Length | 8.69 (7.43-10.1) | 8.76 (7.46-10.18) | 8.56 (7.27-10.03) | 8.41 (7.08-9.8) | 8.57 (7.22-10.16) |
| Lacunar Contrast | 4.8 | 6.2 | 7.8 | 8.8 | 9.5 |
| Signal to Noise | 1.57 | 1.45 | 1.40 | 1.45 | 1.41 |
| Lacuna Number | 3972 | 3743 | 3440 | 3126 | 2676 |
| Lacunar Density | 81.7 | 77.0 | 70.8 | 64.2 | 55.1 |

Data are median (25% - 75% of Inter Quartile Range)

^*^*p*<0.05 compared to ground vehicle, Tukey’s HSD. α set a priori to 0.05.

*Units of Intensity are Arbitrary XRM intensity values

Table 3: Comparison of osteocyte lacunar morphologies and scan durations at different distances between source and detector.

| **Resolution, Objective** | 0.6 µm, 20x | 0.6 µm, 20x |
| --- | --- | --- |
| **Geometric Magnification** | 2.26 | 2.26 |
| **Power and Voltage** | 40kV 3W | 40kV 3W |
| **Source and Detector Distance** | **13.1 mm** | **23.1 mm** |
| **Exposure, Scan Duration** | 9.5 sec, 8.5 hrs | 31.0 sec, 27.6 hrs |
| Lacuna Volume | 232.49 (179.69-314.35) | 263.2 (204.61-345.4) |
| Fitted Ellipsoid Volume | 253.8 (195.69-349.84) | 291.86 (225.94-397.59) |
| % Difference in Volume | -8.07 (-12.5--5.75) | -10.44 (-16.32--7.07) |
| Surface Area | 261.64 (214.55-331.64) | 285.19 (234.76-357.87) |
| Fitted Ellipsoid Surface Area | 258.5 (210.95-326.45) | 287.12 (235.61-357.4) |
| % Difference in Surface Area | 1.69 (-0.33-3.5) | 0.00 (-2-1.77) |
| Volume/Surface Area | 0.89 (0.81-0.96) | 0.91 (0.84-0.99) |
| Lacunar Aspect Ratio | 0.2 (0.15-0.28) | 0.2 (0.14-0.28) |
| Sphericity | 0.47 (0.45-0.50) | 0.48 (0.45-0.51) |
| Lacuna Oblateness | -0.44 (-0.63--0.19) | -0.43 (-0.62--0.18) |
| Lacuna Stretch | 0.79 (0.75-0.82) | 0.8 (0.75-0.82) |
| Lacuna Width | 1.86 (1.64-2.09) | 1.9 (1.7-2.15) |
| Lacuna Height | 3.77 (3.21-4.47) | 3.98 (3.41-4.73) |
| Lacunae Length | 8.69 (7.43-10.1) | 9.22 (7.87-10.67) |
| Lacunar Contrast | 4.8 | 8.1 |
| Signal to Noise | 1.57 | 1.31 |
| Lacunar Number | 3972 | 3954 |
| Lacunar Density | 81.7 | 81.4 |

Data are median (25% - 75% of Inter Quartile Range)

^*^*p*<0.05 compared to ground vehicle, Tukey’s HSD. α set a priori to 0.05.

*Units of Intensity are Arbitrary XRM intensity values

Table 4: Comparison of osteocyte lacunar morphologies in Ground Control (GC) and Microgravity exposure mice at two different anatomical locations in the proximal tibia.

| **Spaceflight Group** | Ground (GC) | Ground (GC) | Microgravity | Microgravity |
| --- | --- | --- | --- | --- |
| **Anatomical Location** | Anterior Medial | Posterior Lateral | Anterior Medial | Posterior Lateral |
| Lacuna Number | 20359 | 15624 | 16703 | 15947 |
| Lacunar Density |  |  |  |  |
| Lacuna Volume | 284.72 (217.26-372.52) | 233.81 (179.62-303.98) | 257.88 (204.05-327.47) | 252.41 (198.32-317.91) |
| Surface Area | 460.65 (372.95-574.76) | 413.58 (338.82-498.47) | 429.32 (359.38-518.86) | 441.07 (367.44-523.37) |
| Volume/Surface Area | 0.61 (0.56-0.67) | 0.56 (0.51-0.62) | 0.6 (0.54-0.65) | 0.57 (0.52-0.63) |
| Lacunar Aspect Ratio | 0.21 (0.15-0.3) | 0.18 (0.13-0.27) | 0.19 (0.14-0.28) | 0.2 (0.14-0.29) |
| Lacuna Oblateness | 0.69 (0.44-0.82) | 0.71 (0.5-0.82) | 0.71 (0.49-0.83) | 0.66 (0.43-0.8) |
| Lacuna Stretch | 0.79 (0.7-0.85) | 0.82 (0.74-0.87) | 0.81 (0.72-0.86) | 0.8 (0.71-0.86) |
| Lacunae Length | 78587.45 (46924.55-131843.83) | 64045.6 (39423.87-99424.87) | 70825.92 (44539.14-108767.55) | 72096.02 (45838.02-108113.65) |

Data are median (25% - 75% of Inter Quartile Range)

^*^*p*<0.05 compared to ground vehicle, Tukey’s HSD. α set a priori to 0.05.

III. ***Figures***

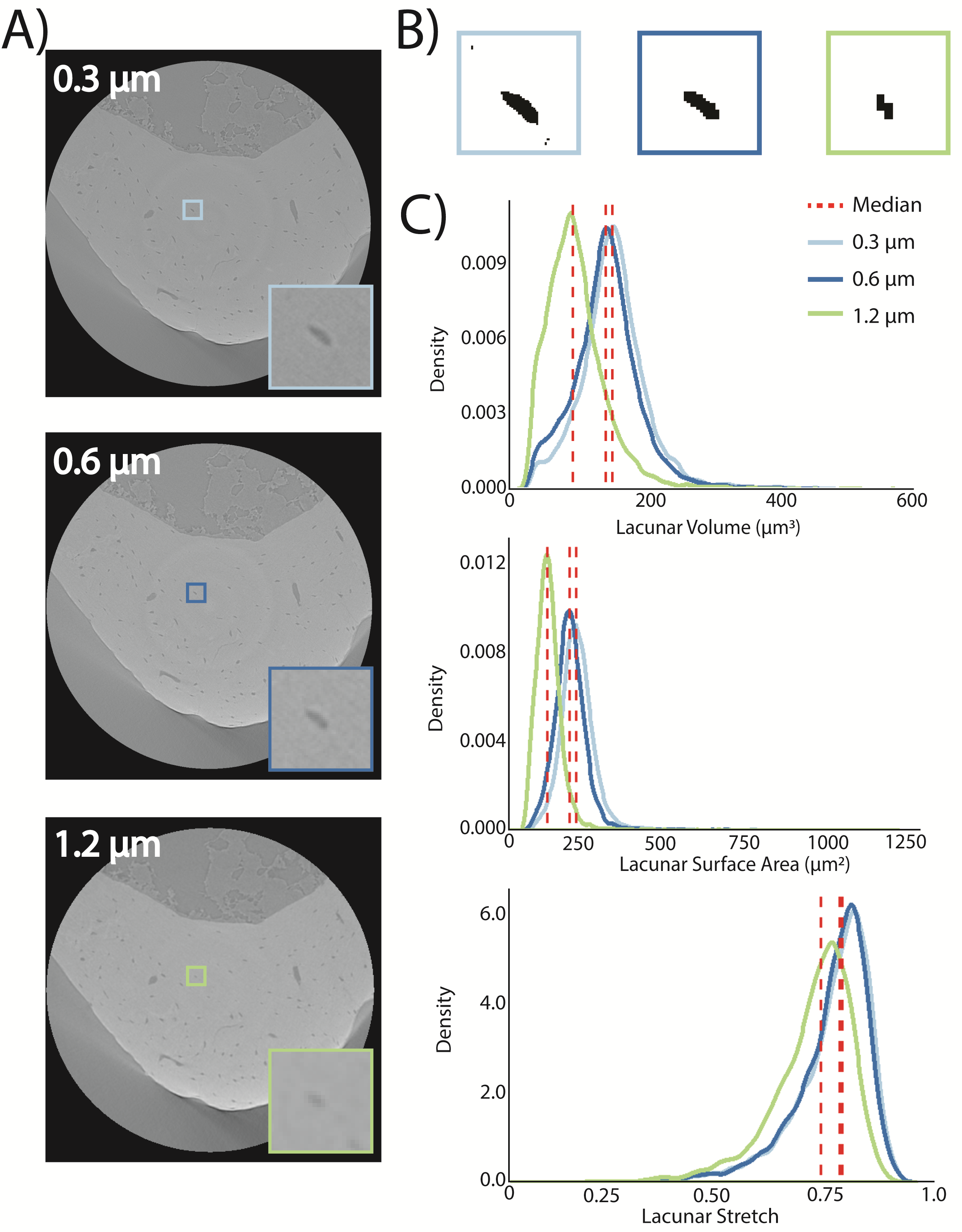

**Supplemental Figure 1. Lower nominal resolution values improve the contrast and segmentation of osteocyte lacunae at the same geometric magnification.** **A)** Posterior region of a mouse tibia was scanned three times using the 20x objective at different resolutions for each scan: 0.3 µm, 0.6 µm, and 1.2 µm resolution. The same osteocyte lacuna is presented at each resolution for comparison as a greyscale. **B)** Comparison of the Otsu algorithm used to binarize the greyscale image for each of the representative osteocyte lacunae from part A). **C)** Distributions of lacunar volume. **D)** Distributions of lacunar surface area. **E)** Distributions of lacunar oblateness. **F)** Distributions of lacunar stretch. Data are presented with a median line (dashed red line) for each of the nominal resolutions’ distributions. Sample sizes, N = 6617 lacunae for 0.3 µm resolution scan, N = 6503 for 0.6 µm resolution scan, and N = 5031 for 1.2 µm resolution scan.

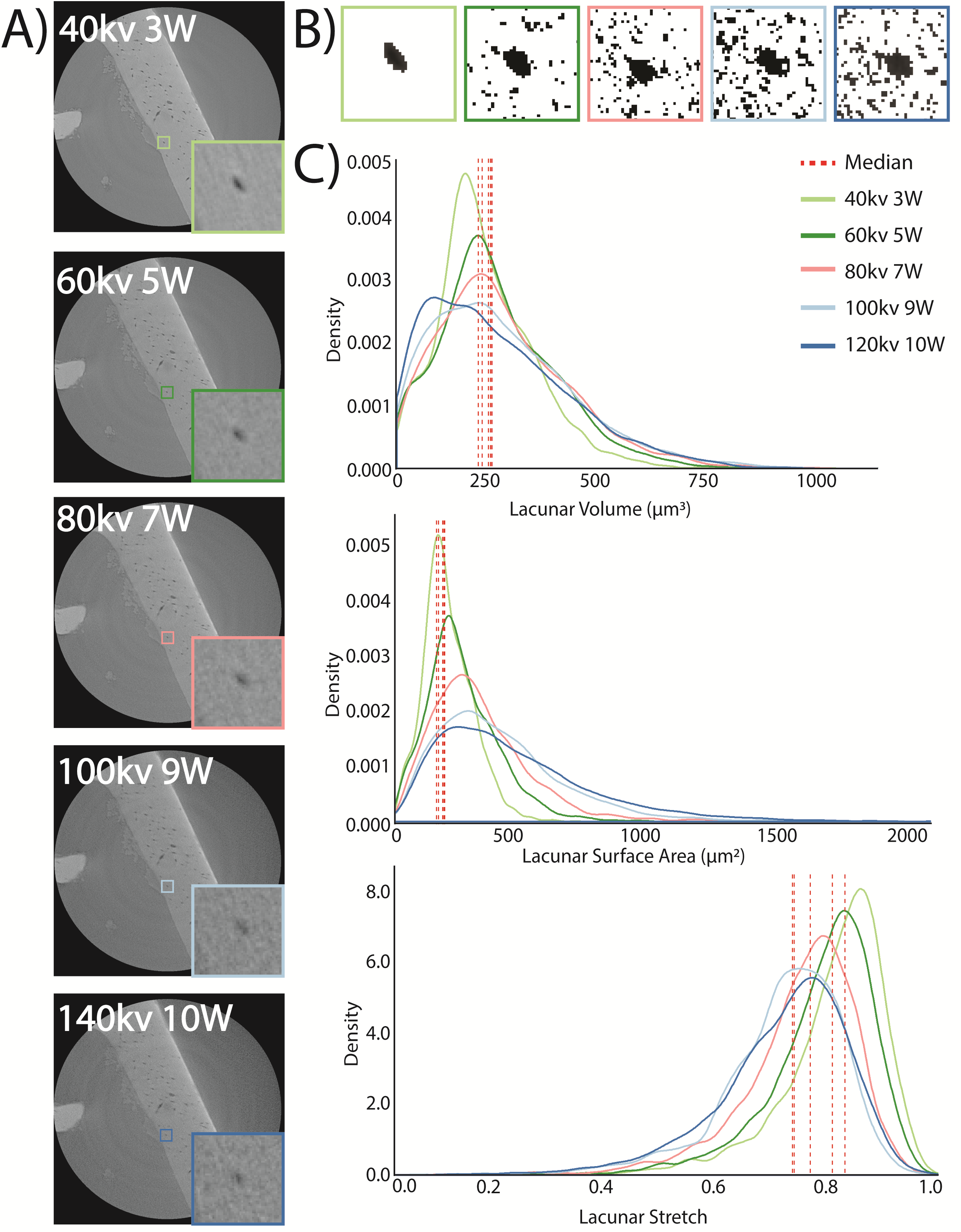

**Supplemental Figure 2. Lower voltage and power values improve the contrast and segmentation of osteocyte lacunae at the same geometric magnification.** **A)** Anterior region of a mouse tibia was scanned five times using the 20x objective with 0.6 µm resolutions for each scan. Power and voltage settings were varied for each scan: 40kv and 3W, 60kv and 5W, 80kv and 7W, 100kv and 9W, 140kv and 10W. The same osteocyte lacuna is presented at each resolution for comparison as a greyscale. **B)** Comparison of the Otsu algorithm used to binarize the greyscale image for each of the representative osteocyte lacunae from part A). **C)** Distributions of lacunar volume. **D)** Distributions of lacunar surface area. **E)** Distributions of lacunar oblateness. **F)** Distributions of lacunar stretch. Data are presented with a median line (dashed red line) for each of the nominal resolutions’ distributions. Sample sizes, N = 3972 lacunae for 40kv and 3W scan, N = 3743 for 60kv and 5W scan, N = 3440 for 80kv and 7W, N = 3126 for 100kv and 9W, N = 2676 for 140kv and 10W.

IV. ***X-Ray Microscopy Parameter Optimization***

A critical need exists for precise and facile assessment of three-dimensional osteocyte lacunar morphometries and standardized reporting of parameters important to consistent image acquisition and analysis. This work focuses on *ex vivo* evaluation of isolated mouse bone specimens using an XRM. Our goal is to provide general guidelines for lacunar morphometry evaluations for a variety of different experimental protocols. These techniques can be used to evaluate changes in the osteocyte’s immediate bone environment, or perilacunar remodeling in future studies of mineral homeostasis, skeletal diseases, and changes in loading environments. While previous work has demonstrated the need for osteocyte lacunar evaluations to be made in three dimensions^(1,2)^, the present study highlighted the importance of instrument parameter selection and sample variability. Collectively, our results demonstrated the importance of reporting parameters that will alter lacunar morphometric measures. To improve future study designs of perilacunar remodeling investigations, standardization of reported parameters for future investigations of osteocyte lacunar morphometries is critically needed. We outline potential analytical steps for evaluating such parameters. Broadly our results demonstrated a need for standardization of reporting imaging parameters and lacunar measurements in the field of orthopaedics so that we will better understand the mechanisms through which osteocytes participate in overall bone health.

***IV.I Geometric Resolution significantly alters osteocyte lacunar morphometry measures.***

To determine the effects of geometric magnification on the precision of osteocyte lacunar morphometries, we scanned the same region of bone using the 4ⅹ and 20x objective with the same nominal resolution of 1.2 µm. The geometric magnification was 2.83 and 2.26 for the 4ⅹ and 20x scans, respectively. Despite having the same nominal resolution, the scan with the lowest geometric magnification resolved the greatest number of lacunae but higher SNR (**Sup.** **Table 1**). With the 4ⅹ objective, 3094 lacunae were resolved whereas, with the 20x objective, 50,314 lacunae were identified in the same sample. Additionally, the 4ⅹ objective scan had a greater median Volume (+179%), Surface Area (+109%), and lower Lacunar Stretch (-21%), and Lacunar Oblateness (-112%) as the 20x objective scan. Both scans required under three hours, but the 20x objective scan had an SNR value of 1.28 compared to 1.10 for the 4ⅹ scan. Similarly, the 20x objective scan had a lower lacunar coefficient of variance (24.0) than the 4ⅹ scan (63.5). Taken together, the 20x objective resulted in scans with osteocyte lacunae with greater contrast to the surround bone matrix which allows for more reliable segmentation and quantification of lacunar morphometries than those visualized with a 4x objective.

***IV.II 0.6 µm scan provides comparable osteocyte lacunar morphometries with one-fourth of the scan time.***

To evaluate the precision of osteocyte lacunar morphometric measures in a time and cost-effective manner, we varied the resolutions for matched scans. Using the 20x objective and/with a constant geometric magnification of 2.26, we varied the nominal resolution (0.3, 0.6, and 1.2 µm) for three separate scans of the same region of bone to determine the effects on lacunar morphometries. For ¼ of the scan time, the 0.6 µm resolution scan produced the same median values of all the lacunar parameters as did the 0.3 µm scan. All three resolutions had the same median values of lacunar stretch, oblateness, height, and width, but the 0.3 µm scan had a smaller median Volume (-35%) and Surface Area (-38%) than the 1.2 µm scan **(Sup. Fig. 1C., Sup. Table 1)**. Despite the similarities, the 0.3 µm resolution scan required 35.6 hours of scan time and resolved 6617 lacunae, whereas the 0.6 µm resolution scan needed only 8.5 hours for identification of 6503 lacunae, but only 50,314 lacunae at 1.2 µm resolution in 2.9 hours. SNR for all of the scans was comparable - 1.28 for the 0.3 and 1.2 µm resolution scans, and 1.27 for the 0.6 µm resolution scan. By contrast, lacunar coefficient of variance was inversely proportional to scan resolution; the 0.3 µm scan had the lowest value (12.8), followed by the 0.6 µm scan (14.5), and the 1.2 µm resolution scan (24.0). As a result, the lacunae of the 0.3 µm and 0.6 µm resolution scans more were more reliably binarized using the Otsu method than the 1.2 µm resolution scan **(Sup. Fig. 1B.)**. Given the shorter scan time required for the 0.6 µm scan, we utilized those parameters (i.e., 20x objective, and 0.6 µm nominal resolution) for subsequent scans examining the impacts of microgravity.

***IV.III Increased power and voltage decreased time and Signal to Noise, but not Lacunar Volume or Oblateness.***

To test if power and voltage affected the precision of lacunar morphometries, we scanned the same region of bone from previously described scans with matched geometric magnification (2.26, objective of 20x), and nominal resolution (0.6 µm resolution). We then varied power and voltage settings using five different settings - power and voltages of: 40 kV/3W, 60 kV/5W, 80 kV/7W, 100 kV/9W, and 140 kV/10W (**Sup.** **Fig. 2A, Sup. Table 2**). The lowest power and voltage parameters had the highest SNR and resolved the greatest number of osteocyte lacunae, but also had the longest scan duration. All scans had the same median value for Volume, Oblateness, and Length, yet 40kV/3W had a comparable median value of Surface Area, and Lacuna Stretch to the 60kV/5W scan (**Sup. Fig. 2C, Sup. Table 2**). For the remaining scans, 3972 lacunae from 40kV/3W, 3743 from 60kV/5W, 3440 from 80kV/7W, 3126 from 100kV/9W, and 2676 from 140kV/100W were resolved. The 40kV/3W scan lasted less than 8.5 hours but had the highest SNR value of 1.57, whereas all other scans were completed under three hours with SNR values ranging 1.40 - 1.45. Lacunar Coefficient of Variance was lowest for 40kV (4.8) and increased with higher powers and voltages. For example, 60kV lacunar coefficient of variance was 30% higher and 140kV was 98% higher. As with resolution, these differences in SNR and Lacunar Coefficient of Variance resulted in varied reliability in binarization of the lacunae (**Sup.** **Fig. 2B.**). Thus, the power and voltage setting of 40kV/3W maximized contrast between lacunae and the bone matrix while minimizing scan duration, and were used with subsequent scans examining the impacts of microgravity.

***IV.IV Reduced distance between source and detector improves the signal to noise and scan duration.***

Next, we tested how the distance between source and detector affected the precision of lacunar morphometries. Using the previously-determined parameters (2.26 geometric magnification, objective of 20x, 0.6 µm nominal resolution, 40 kV/3W voltage and power), we varied the source and detector distance by 10 mm for two scans (13.1 and 23.1 mm). While all the osteocyte lacunar morphometrics had the same median volume values and the number of lacunae were comparable, the scan with the shorter source and detector distance took less than a third of the time and with a higher SNR. Both scans had the same median value of lacuna volume, surface area, oblateness, stretch, and length with n = 3972 lacunae imaged at 13 mm distance and 3954 lacunae at 23 mm between the source and detector (**Supp Table 3**). However, with 23 mm between the source and detector, there was a greater interquartile range for Volume and Surface Area, and a lower SNR of 13.1, while the SNR value for the 13 mm scan was 1.57. Additionally, the 23 mm scan required 27.6 hours whereas the 13 mm scan lasted less than 8.5 hours. Lacunar coefficient of variance was 14.2 for the closer scan, but 8.1 for the more distant scan. However, the scan with greater source and detector distance had an 18% greater mean intensity per lacunae, and 108% greater standard deviation of intensity per lacunae as compared to the closer scan. For subsequent scans, we thus utilized the shortest source-detector distance. For subsequent scans, we thus utilized the shortest source-detector distance to minimize instrument time and maximize contrast between lacunae and the bone matrix.

***IV.V Suggested Parameters***

This study evaluated the tradeoffs that are needed to maximize data quality while reducing scan time and cost. Here, we found that geometric magnification, nominal resolution, power and voltage, and distance between source and detector all affect quantitative measures of osteocyte lacunar morphometry as well as scan duration. 0.3 µm resolution scans provided the highest signal-to-noise ratios along with the lowest lacunar coefficient of variance, the greatest number of lacunae, and the lowest percent error on volume and surface area, but they were not cost-effective. At this resolution, scans required long imaging times (> 20 hours) and may increase the risk of failure and sample drift. By contrast, 0.6 µm scans were ideal for optimal through-put sample evaluation, as they produced similar lacunar morphometries measures and required one-tenth of the total time required by the 0.3 µm resolution scan. Additionally, the selection of nominal resolution should be consistent for any comparison of lacunar morphometries and reflective of the research questions. Surface area was most varied by resolution and should only be compared between samples with the same resolution. Additionally, osteocyte lacunae are not perfect ellipses, though this common assumption may be sufficient for coarser resolutions (1.2 µm) than finer ones (0.3 µm). Ultimately, we determined using a higher geometric magnification can dramatically reduce the scan duration but compromise the reliability of the data.

Like geometric magnification, higher power and voltage can shorten the duration of the scan but at the expense of reliable lacunar segmentation and identification. Higher voltage and power can distort the shape of the lacunae, or other features in the bone, as it reduces the contrast between the lacunae and the surrounding bone, decreasing the signal-to-noise ratio. With decreased contrast, some lacunae cannot be detected or segmented in the post-processing of the scans. Alternatively, lower power and voltage provide better contrast between the bone and air or liquid-filled cavities within it. The use of a synchrotron source, which uses lower power and voltage ranges than most commercial X-Ray microscopes can achieve, can evaluate the contrasts in bone mineral density using phase contrast^(3)^. Leveraging high resolution with lower power and voltage parameters may allow for future investigations of how osteocytes modulate bone mineral density in the peri-lacunar bone during development or shifts in mineral homeostasis.

While our results suggest ideal parameters to improve the visualization and quantification of osteocyte lacunae in mouse bones using a Zeiss Xradia Versa, we seek to demonstrate the importance of reporting specific imaging parameters when using either X-ray microscopy or µCT for osteocyte lacunae. To improve the repeatability and reliability of future studies of lacunar morphometries, we suggest researchers report geometric magnification, nominal resolution, power and voltage, exposure time, and source and detector distance (either from each other or from the sample). Additionally, the measurements of lacunae also need to be standardized to help make comparisons between research teams and study designs. To take the first step in standardizing measures of lacunar morphologies we have made our custom python code open source through Dragonfly Infinite Toolbox. This plugin will allow researchers to compute lacunar oblateness, stretch, length, width, height, fitted ellipsoidal volume, and fitted ellipsoidal surface area on any sample. Furthermore, measures of lacunar density need to be defined consistently. Currently, there are several definitions including Lacuna volume Fraction Lc.V/TV (%)^(4)^ Lacuna density N.Lc/TV (1/mm^3^)^(4)^ We suggest the intuitive and meaningful measure of Lc. N/BV (mm^3^)^(5)^. With consistent and standardized measurement reporting across the orthopaedic community, we will be able to compare changes in osteocyte lacunar morphologies and better understand the role of PLR in aging, disuse, and disease.

V. ***Python Plug-In for Dragonfly***

Custom code (Python) was developed to evaluate lacunar morphometry measures of lacunar oblateness, stretch, length, height and width, and implemented in Dragonfly Pro to analyze the shape of individual lacunae within each sample. This code is available through The Objects Research Systems (ORS) Infinite Toolbox at the following link and as a part of this document.

Dragonfly Infinite Toolbox: <https://infinitetoolbox.theobjects.com/>

Analyze Osteocyte Lacunae Shapes

:author: Jenn Coulombe

:contact:

:

:organization: CU Boulder

:address: 1111 Engineering Drive, Boulder CO, 80309

:copyright:

:date: Mar 23 2018 13:00

:dragonflyVersion: 3.1.0.319

:UUID: 6d6b028a2ecc11e881c1509a4c54acff

"""

__version__ = '1.0.0'

import numpy as np

import math

from ORSModel import orsObj, DimensionUnit

import COMWrapper.ORS_def as ORSDef

from OrsHelpers.arrayhelper import ArrayHelper

from OrsPythonPlugins.OrsObjectAnalysis.PythonScriptsStatisticsGenerators.StatisticsGeneratorAbstract import StatisticsGeneratorAbstract

from OrsLibraries.preferences import Preferences

from COMWrapper.ORS_def import CxvUniverse_Dimension

class ShapeAnalyzer_6d6b028a2ecc11e881c1509a4c54acff(StatisticsGeneratorAbstract):

"""

Methods to overload are described in the class StatisticsGeneratorAbstract

(file: ORSPYTHON\OrsPythonPlugins\OrsObjectAnalysis\PythonScriptsStatisticsGenerators\StatisticsGeneratorAbstract.py)

See also: class StatisticsGeneratorBasic

(file: ORSPYTHON\OrsPythonPlugins\OrsObjectAnalysis\PythonScriptsStatisticsGenerators\StatisticsGeneratorBasic.py)

"""

def __init__(self):

super().__init__()

self.outputTagsNoDataset = ["Lacuna Stretch ", "Lacuna Oblateness ", "Primary Orientation Theta ", "Primary Orientation Phi ",

"Secondary Orientation Theta ", "Secondary Orientation Phi ", "Lacuna Length ", "Lacuna Height ", "Lacuna Width ",

"Span Theta ", "Fitted Ellipsoid Volume ", "Fitted Ellipsoid Surface Area "]

#Example: self.outputTagsNoDataset = ['Sphericity']

self.outputTagsWithDataset = [] #Put here the tags for computation where a dataset is required

@classmethod

def getUUID(cls):

return "6d6b028a2ecc11e881c1509a4c54acff"

def getOutputTagsWithoutDataset(self):

return self.outputTagsNoDataset

def getOutputTagsWithDataset(self):

return self.outputTagsWithDataset

def generateOutputsWithoutDataset(self, multiROIGUID, listTagsRequested=None, IProgress=None):

multi_roi = orsObj(multiROIGUID) #Get the multi roi object from his guid

t_step = 0

#Compute a numpy array of all multi ROI non empty labels

arrayNonEmptyLabels = multi_roi.getNonEmptyLabels(None)

npArrayNonEmptyLabels = ArrayHelper.ConvertOrsToNumpyArray(arrayNonEmptyLabels)

arrayNonEmptyLabels.deleteObject()

#Here is an integrated cache mechanic. The function readScalarValues check if the statistical properties had

#already been previously computed. If so, it fills the npArrayDictToReturn dictionary with the computed values

#and remove the corresponding statistical properties tag from listTagsRemaining

listTagsRemaining = listTagsRequested

if listTagsRemaining is None:

listTagsRemaining = self.getOutputTagsWithoutDataset()

npArrayDictToReturn = self.readScalarValues(multi_roi, npArrayNonEmptyLabels, listTagsRemaining)

stats = 0

if any(x in ["Lacuna Stretch ", "Lacuna Oblateness ", "Primary Orientation Theta ", "Primary Orientation Phi ",

"Secondary Orientation Theta ", "Secondary Orientation Phi ", "Lacuna Length ", "Lacuna Height ",

"Lacuna Width ", "Span Theta ", "Fitted Ellipsoid Volume ",

"Fitted Ellipsoid Surface Area "] for x in listTagsRemaining):

stats |= ORSDef.CXV_LABELED_MULTI_ROI_STATS_TENSOR_INERTIA

if stats != 0: #Validated for 3.1

labeled_multi_roi_analyzer = multi_roi.generateAnalyzer(None, None, t_step, stats, False, IProgress)

#At last, we have to compute statistical properties that hadn't been previously computed

non_empty_label_count = len(npArrayNonEmptyLabels)

for tag in listTagsRemaining: #iterate on remaining tags

if IProgress is not None and IProgress.getIsCancelled(): #Check if user cancel the computation

#If user cancel the computation, return zeros

npArrayDictToReturn[tag] = (np.zeros((non_empty_label_count,), dtype=int), True)

elif tag == "Lacuna Stretch ":

#(Largest - Smallest) / Largest

npArrayResult = np.zeros(non_empty_label_count, dtype=float)

for arrayIndex, labelIndex in enumerate(npArrayNonEmptyLabels):

#Get all eigenvalues

smallestEigenValue = labeled_multi_roi_analyzer.getLabelInertiaEigenValue(labelIndex, 0)

intermediateEigenValue = labeled_multi_roi_analyzer.getLabelInertiaEigenValue(labelIndex, 1)

largestEigenValue = labeled_multi_roi_analyzer.getLabelInertiaEigenValue(labelIndex, 2)

#Get voxel fill volume

voxelCount = multi_roi.getLabelSize(labelIndex)

#Scale factor between voxels and meters

spacingX = multi_roi.getXSpacing()

#Calculate radii

smallestRadius = (math.pow(2.5*(smallestEigenValue+intermediateEigenValue-largestEigenValue)

/voxelCount,0.5))*spacingX

largestRadius = (math.pow(2.5*(intermediateEigenValue+largestEigenValue-smallestEigenValue)

/voxelCount,0.5))*spacingX

#Calculate stretch

lacunastretch = ((largestRadius-smallestRadius)/largestRadius)

npArrayResult[arrayIndex] = lacunastretch

npArrayDictToReturn[tag] = (npArrayResult, False)

elif tag == "Lacuna Oblateness ":

#(2 * ((Intermediate - Smallest)/(Largest - Smallest))) - 1

npArrayResult = np.zeros(non_empty_label_count, dtype=float)

for arrayIndex, labelIndex in enumerate(npArrayNonEmptyLabels):

#Get all eigenvalues

smallestEigenValue = labeled_multi_roi_analyzer.getLabelInertiaEigenValue(labelIndex, 0)

intermediateEigenValue = labeled_multi_roi_analyzer.getLabelInertiaEigenValue(labelIndex, 1)

largestEigenValue = labeled_multi_roi_analyzer.getLabelInertiaEigenValue(labelIndex, 2)

#Get voxel fill volume

voxelCount = multi_roi.getLabelSize(labelIndex)

#Scale factor between voxels and meters

spacingX = multi_roi.getXSpacing()

#Calculate radii

smallestRadius = (math.pow(2.5 * (smallestEigenValue + intermediateEigenValue - largestEigenValue)

/ voxelCount, 0.5)) * spacingX

intermediateRadius = (math.pow(2.5*(smallestEigenValue+largestEigenValue-intermediateEigenValue)

/voxelCount,0.5))*spacingX

largestRadius = (math.pow(2.5*(intermediateEigenValue+largestEigenValue-smallestEigenValue)

/voxelCount,0.5))*spacingX

#Calculate oblateness

lacunaOblateness = (2*((intermediateRadius-smallestRadius)/(largestRadius-smallestRadius)))-1

npArrayResult[arrayIndex] = lacunaOblateness

npArrayDictToReturn[tag] = (npArrayResult, False)

elif tag == "Lacuna Width ":

#The absolute value of the smallest radius

npArrayResult = np.zeros(non_empty_label_count, dtype=float)

for arrayIndex, labelIndex in enumerate(npArrayNonEmptyLabels):

#Get all eigenvalues

smallestEigenValue = labeled_multi_roi_analyzer.getLabelInertiaEigenValue(labelIndex, 0)

intermediateEigenValue = labeled_multi_roi_analyzer.getLabelInertiaEigenValue(labelIndex, 1)

largestEigenValue = labeled_multi_roi_analyzer.getLabelInertiaEigenValue(labelIndex, 2)

#Get voxel fill volume

voxelCount = multi_roi.getLabelSize(labelIndex)

#Scale factor between voxels and meters

spacingX = multi_roi.getXSpacing()

#Calculate radius

smallestRadius = math.pow(2.5*(smallestEigenValue+intermediateEigenValue-largestEigenValue)

/voxelCount,0.5)*spacingX

npArrayResult[arrayIndex] = smallestRadius

npArrayDictToReturn[tag] = (npArrayResult, False)

elif tag == "Lacuna Height ":

#The absolute value of the intermediate radius

npArrayResult = np.zeros(non_empty_label_count, dtype=float)

for arrayIndex, labelIndex in enumerate(npArrayNonEmptyLabels):

#Get all eigenvalues

smallestEigenValue = labeled_multi_roi_analyzer.getLabelInertiaEigenValue(labelIndex, 0)

intermediateEigenValue = labeled_multi_roi_analyzer.getLabelInertiaEigenValue(labelIndex, 1)

largestEigenValue = labeled_multi_roi_analyzer.getLabelInertiaEigenValue(labelIndex, 2)

#Get voxel fill volume

voxelCount = multi_roi.getLabelSize(labelIndex)

#Scale factor between voxels and meters

spacingX = multi_roi.getXSpacing()

#Calculate radius

intermediateRadius = math.pow(2.5*(smallestEigenValue+largestEigenValue-intermediateEigenValue)

/voxelCount,0.5)*spacingX

npArrayResult[arrayIndex] = intermediateRadius

npArrayDictToReturn[tag] = (npArrayResult, False)

elif tag == "Lacuna Length ":

#The absolute value of the largest radius

npArrayResult = np.zeros(non_empty_label_count, dtype=float)

for arrayIndex, labelIndex in enumerate(npArrayNonEmptyLabels):

#Get all eigenvalues

smallestEigenValue = labeled_multi_roi_analyzer.getLabelInertiaEigenValue(labelIndex, 0)

intermediateEigenValue = labeled_multi_roi_analyzer.getLabelInertiaEigenValue(labelIndex, 1)

largestEigenValue = labeled_multi_roi_analyzer.getLabelInertiaEigenValue(labelIndex, 2)

#Get voxel fill volume

voxelCount = multi_roi.getLabelSize(labelIndex)

#Scale factor between voxels and meters

spacingX = multi_roi.getXSpacing()

#Calculate radius

largestRadius = math.pow(2.5*(largestEigenValue+intermediateEigenValue-smallestEigenValue)

/voxelCount,0.5)*spacingX

npArrayResult[arrayIndex] = largestRadius

npArrayDictToReturn[tag] = (npArrayResult, False)

elif tag == "Primary Orientation Theta ":

#Largest eigenvector

npArrayResult = np.zeros(non_empty_label_count, dtype=float)

for arrayIndex, labelIndex in enumerate(npArrayNonEmptyLabels):

largestEigenVector = labeled_multi_roi_analyzer.getLabelInertiaEigenVector(labelIndex, 2)

largestEigenVectorNorm = largestEigenVector.getNormalized()

#worldTransform = multi_roi.getBox().getWorldTranformation()

#largestEigenVecWorld = worldTransform.getTransformedVector(largestEigenVectorNorm)

#PrimOrient = largestEigenVecWorld.getTheta()

PrimOrientTheta = largestEigenVectorNorm.getTheta()

npArrayResult[arrayIndex] = PrimOrientTheta

npArrayDictToReturn[tag] = (npArrayResult, False)

elif tag == "Primary Orientation Phi ":

#Largest eigenvector

npArrayResult = np.zeros(non_empty_label_count, dtype=float)

for arrayIndex, labelIndex in enumerate(npArrayNonEmptyLabels):

largestEigenVector = labeled_multi_roi_analyzer.getLabelInertiaEigenVector(labelIndex, 2)

largestEigenVectorNorm = largestEigenVector.getNormalized()

#worldTransform = multi_roi.getBox().getWorldTranformation()

#largestEigenVecWorld = worldTransform.getTransformedVector(largestEigenVectorNorm)

#PrimOrient = largestEigenVecWorld.getTheta()

PrimOrientPhi = largestEigenVectorNorm.getPhi()

npArrayResult[arrayIndex] = PrimOrientPhi

npArrayDictToReturn[tag] = (npArrayResult, False)

elif tag == "Secondary Orientation Theta ":

#Intermediate eigenvector

npArrayResult = np.zeros(non_empty_label_count, dtype=float)

for arrayIndex, labelIndex in enumerate(npArrayNonEmptyLabels):

intermediateEigenVector = labeled_multi_roi_analyzer.getLabelInertiaEigenVector(labelIndex, 1)

intermediateEigenVectorNorm = intermediateEigenVector.getNormalized()

SecondOrientTheta = intermediateEigenVectorNorm.getTheta()

npArrayResult[arrayIndex] = SecondOrientTheta

npArrayDictToReturn[tag] = (npArrayResult, False)

elif tag == "Secondary Orientation Phi ":

#Intermediate eigenvector

npArrayResult = np.zeros(non_empty_label_count, dtype=float)

for arrayIndex, labelIndex in enumerate(npArrayNonEmptyLabels):

intermediateEigenVector = labeled_multi_roi_analyzer.getLabelInertiaEigenVector(labelIndex, 1)

intermediateEigenVectorNorm = intermediateEigenVector.getNormalized()

SecondOrientPhi = intermediateEigenVectorNorm.getPhi()

npArrayResult[arrayIndex] = SecondOrientPhi

npArrayDictToReturn[tag] = (npArrayResult, False)

elif tag == "Span Theta ":

#deviation from average orientation

npArrayResult = np.zeros(non_empty_label_count, dtype=float)

Orientsum = 0

counter = 0

for arrayIndex, labelIndex in enumerate(npArrayNonEmptyLabels):

largestEigenVector = labeled_multi_roi_analyzer.getLabelInertiaEigenVector(labelIndex, 2)

PrimOrient = largestEigenVector.getTheta()

Orientsum = Orientsum + PrimOrient

counter = counter + 1

AvgOrient = Orientsum/counter

for arrayIndex, labelIndex in enumerate(npArrayNonEmptyLabels):

largestEigenVector = labeled_multi_roi_analyzer.getLabelInertiaEigenVector(labelIndex, 2)

PrimOrient = largestEigenVector.getTheta()

Deviation = AvgOrient - PrimOrient

npArrayResult[arrayIndex] = Deviation

npArrayDictToReturn[tag] = (npArrayResult, False)

#Fitted Ellipsoid Calculations

elif tag == "Fitted Ellipsoid Volume ":

npArrayResult = np.zeros(non_empty_label_count, dtype=float)

for arrayIndex, labelIndex in enumerate(npArrayNonEmptyLabels):

#Get all eigenvalues

smallestEigenValue = labeled_multi_roi_analyzer.getLabelInertiaEigenValue(labelIndex, 0)

intermediateEigenValue = labeled_multi_roi_analyzer.getLabelInertiaEigenValue(labelIndex, 1)

largestEigenValue = labeled_multi_roi_analyzer.getLabelInertiaEigenValue(labelIndex, 2)

#Get voxel fill volume

voxelCount = multi_roi.getLabelSize(labelIndex)

#Scale factor between voxels and meters

spacingX = multi_roi.getXSpacing()

#Calculate radii

smallestRadius = (math.pow(2.5*(intermediateEigenValue+largestEigenValue-smallestEigenValue)

/voxelCount,0.5))*spacingX

intermediateRadius = (math.pow(2.5*(smallestEigenValue+largestEigenValue-intermediateEigenValue)

/voxelCount,0.5))*spacingX

largestRadius = (math.pow(2.5*(smallestEigenValue+intermediateEigenValue-largestEigenValue)

/voxelCount,0.5))*spacingX

#Calculate volume

FittedEllipsoidVol = (4/3)*(math.pi)*largestRadius*intermediateRadius*smallestRadius

npArrayResult[arrayIndex] = FittedEllipsoidVol

npArrayDictToReturn[tag] = (npArrayResult, False)

elif tag == "Fitted Ellipsoid Surface Area ":

npArrayResult = np.zeros(non_empty_label_count, dtype=float)

for arrayIndex, labelIndex in enumerate(npArrayNonEmptyLabels):

#Get all eigenvalues

smallestEigenValue = labeled_multi_roi_analyzer.getLabelInertiaEigenValue(labelIndex, 0)

intermediateEigenValue = labeled_multi_roi_analyzer.getLabelInertiaEigenValue(labelIndex, 1)

largestEigenValue = labeled_multi_roi_analyzer.getLabelInertiaEigenValue(labelIndex, 2)

#Get voxel fill volume

voxelCount = multi_roi.getLabelSize(labelIndex)

#Scale factor between voxels and meters

spacingX = multi_roi.getXSpacing()

#Calculate radii

smallestRadius = (math.pow(2.5*(intermediateEigenValue+largestEigenValue-smallestEigenValue)

/voxelCount,0.5))*spacingX

intermediateRadius = (math.pow(2.5*(smallestEigenValue+largestEigenValue-intermediateEigenValue)

/voxelCount,0.5))*spacingX

largestRadius = (math.pow(2.5*(smallestEigenValue+intermediateEigenValue-largestEigenValue)

/voxelCount,0.5))*spacingX

#Calculate SA

FittedEllipsoidSA = 4*(math.pi)*(math.pow(((math.pow(largestRadius*intermediateRadius,1.6))+

(math.pow(smallestRadius*intermediateRadius,1.6))+

(math.pow(largestRadius*smallestRadius,1.6)))/3,(1/1.6)))

npArrayResult[arrayIndex] = FittedEllipsoidSA

npArrayDictToReturn[tag] = (npArrayResult, False)

\# Put here an "elif" for each supported tag of self.outputTagsNoDataset

\# Example:

\# elif tag == "Sphericity":

\#	import math

\#	from ORSModel.ors import MultiROIAnalyzer

\#	# Compute stats using an instance of MultiRoiAnalyzer, an Ors object used to compute basic statistics on

\#	# multi ROI

\#	multi_roi_analyzer = multi_roi.generateAnalyzer(None, None, t_step,

\#	MultiROIAnalyzer.stats.surface_area,

\#	False, IProgress)

\#

\#	# First, compute volume

\#	labels_volume = multi_roi_analyzer.getIndiciesCountInLabels() # Voxel count per label (spacing is not considered)

\#	np_labels_volume = ArrayHelper.ConvertOrsToNumpyArray(labels_volume) # Convert to numpy array

\#	np_labels_volume_non_empty_labels = np_labels_volume[npArrayNonEmptyLabels] # Remove empty label

\#

\#	# convert to meter

\#	volume_per_voxel_in_meters = multi_roi.getXSpacing() * multi_roi.getYSpacing() * multi_roi.getZSpacing()

\#	np_labels_volume_non_empty_labels = np.multiply(np_labels_volume_non_empty_labels, volume_per_voxel_in_meters)

\#

\#	# Then compute surface area

\#	labels_surface_area = multi_roi_analyzer.getLabelsSurfaceArea() # Surface area considering spacing

\#	np_labels_surface_area = ArrayHelper.ConvertOrsToNumpyArray(labels_surface_area) # Convert to numpy array

\#	np_labels_surface_area_non_empty_labels = np_labels_surface_area[npArrayNonEmptyLabels] # Remove empty label

\#

\#	# Compute Sphericity https://en.wikipedia.org/wiki/Sphericity

\#	np_labels_sphericity_non_empty_labels = (math.pow(math.pi, 1.0 / 3.0) * np.power((6 * np_labels_volume_non_empty_labels), (2.0 / 3.0))) / np_labels_surface_area_non_empty_labels

\#

\#	# Store value in npArrayDictToReturn

\#	npArrayDictToReturn[tag] = (np_labels_sphericity_non_empty_labels, False)

else:

npArrayDictToReturn[tag] = (np.zeros((non_empty_label_count,), dtype=int), True) # Tag not found

return npArrayDictToReturn

def generateOutputsWithDataset(self, datasetGUID, multiROIGUID, listTagsRequested=None, IProgress=None):

multi_roi = orsObj(multiROIGUID) #Get the multi roi object from his guid

t_step = 0

#Compute a numpy array of all multi ROI non empty labels

arrayNonEmptyLabels = multi_roi.getNonEmptyLabels(None)

npArrayNonEmptyLabels = ArrayHelper.ConvertOrsToNumpyArray(arrayNonEmptyLabels)

arrayNonEmptyLabels.deleteObject()

#Here is an integrated cache mechanic. The function readScalarValues check if the statistical properties had

#already been previously computed. If so, it fills the npArrayDictToReturn dictionary with the computed values

#and remove the corresponding statistical properties tag from listTagsRemaining

listTagsRemaining = listTagsRequested

if listTagsRemaining is None:

listTagsRemaining = self.getOutputTagsWithDataset()

npArrayDictToReturn = self.readScalarValues(multi_roi, npArrayNonEmptyLabels, listTagsRemaining)

#At last, we have to compute statistical properties that hadn't been previously computed

non_empty_label_count = len(npArrayNonEmptyLabels)

for tag in listTagsRemaining: #iterate on remaining tags

if IProgress is not None and IProgress.getIsCancelled(): #Check if user cancel the computation

#If user cancel the computation, return zeros

npArrayDictToReturn[tag] = (np.zeros((non_empty_label_count,), dtype=int), True)

#Put here an "elif" for each supported tag of self.outputTagsWithDataset

else:

npArrayDictToReturn[tag] = (np.zeros((non_empty_label_count,), dtype=int), True) #Tag not found

return npArrayDictToReturn

def updateDefaultDimensionUnit(self, outputTag, view):

pass

#Put here a switch on "outputTag" for each supported tag of self.outputTagsNoDataset and self.outputTagsWithDataset

#Example, for a volumic statistic (the value should be returned in npArrayDictToReturn in the unit of m^3):

#if outputTag == "MyVolume":

\#	self.defaultDimensionUnitDict[outputTag] = view.getVolumeDimensionUnit()

#Example, for a surfacic statistic (the value should be returned in npArrayDictToReturn in the unit of m^2):

#elif outputTag == "MySurface":

\#	self.defaultDimensionUnitDict[outputTag] = view.getSurfaceDimensionUnit()

if outputTag in ['Lacuna Length', 'Lacuna Height', 'Lacuna Width',

'Fitted Ellipsoid Volume', 'Fitted Ellipsoid Surface Area']:

self.defaultDimensionUnitDict[outputTag] = view.getLengthDimensionUnit()

def getDefaultDimensionUnitDict(self):

if self.defaultDimensionUnitDict is None:

self.defaultDimensionUnitDict = {} # Initialization

#Put here each supported tag of self.outputTagsNoDataset and self.outputTagsWithDataset

#with their respective default unit object

#Example:

#self.defaultDimensionUnitDict['MyVolume'] = Preferences.getDefaultVolumeUnit()

#self.defaultDimensionUnitDict['MySurface'] = Preferences.getDefaultSurfaceUnit()

self.defaultDimensionUnitDict['Lacuna Length'] = Preferences.getDefaultLengthUnit()

self.defaultDimensionUnitDict['Lacuna Height'] = Preferences.getDefaultLengthUnit()

self.defaultDimensionUnitDict['Lacuna Width'] = Preferences.getDefaultLengthUnit()

self.defaultDimensionUnitDict['Fitted Ellipsoid Volume'] = Preferences.getDefaultVolumeUnit()

self.defaultDimensionUnitDict['Fitted Ellipsoid Surface Area'] = Preferences.getDefaultSurfaceUnit()

return self.defaultDimensionUnitDict

def getDataDimensionUnitDict(self):

if self.dataDimensionUnitDict is None:

unitFactory = DimensionUnit()

self.dataDimensionUnitDict = {} # Initialization

#Put here each supported tag of self.outputTagsNoDataset and self.outputTagsWithDataset

#with their respective data dimension unit

#Example:

#self.dataDimensionUnitDict["MyVolume"] = unitFactory.getUnitForID(CxvUniverse_Dimension.CXV_DIMENSION_METER_VOLUME)

#self.dataDimensionUnitDict["MySurface"] = unitFactory.getUnitForID(CxvUniverse_Dimension.CXV_DIMENSION_METER_SURFACE)

self.dataDimensionUnitDict['Lacuna Length'] = unitFactory.getUnitForID(

CxvUniverse_Dimension.CXV_DIMENSION_METER_LENGTH)

self.dataDimensionUnitDict['Lacuna Height'] = unitFactory.getUnitForID(

CxvUniverse_Dimension.CXV_DIMENSION_METER_LENGTH)

self.dataDimensionUnitDict['Lacuna Width'] = unitFactory.getUnitForID(

CxvUniverse_Dimension.CXV_DIMENSION_METER_LENGTH)

self.dataDimensionUnitDict['Fitted Ellipsoid Volume'] = unitFactory.getUnitForID(

CxvUniverse_Dimension.CXV_DIMENSION_METER_VOLUME)

self.dataDimensionUnitDict['Fitted Ellipsoid Surface Area'] = unitFactory.getUnitForID(

CxvUniverse_Dimension.CXV_DIMENSION_METER_SURFACE)

unitFactory.deleteObject()

return self.dataDimensionUnitDict

VI. ***References***

1.	Heveran CM, Rauff A, King KB, Carpenter RD, Ferguson VL. A new open-source tool for measuring 3D osteocyte lacunar geometries from confocal laser scanning microscopy reveals age-related changes to lacunar size and shape in cortical mouse bone. Bone. 2018;110:115–27.

2.	Yee CS, Schurman CA, White CR, Alliston T. Investigating Osteocytic Perilacunar/Canalicular Remodeling. Curr Osteoporos Rep. 2019 Jun 22;

3.	Hesse B, Varga P, Langer M, Pacureanu A, Schrof S, Männicke N, et al. Canalicular network morphology is the major determinant of the spatial distribution of mass density in human bone tissue: Evidence by means of synchrotron radiation phase-contrast nano-CT. J Bone Miner Res. 2015;30(2):346–56.

4.	Gerbaix M, Gnyubkin V, Farlay D, Olivier C, Ammann P, Courbon G, et al. One-month spaceflight compromises the bone microstructure, tissue-level mechanical properties, osteocyte survival and lacunae volume in mature mice skeletons. Sci Rep. 2017 Dec 1;7(1):2659.

5.	Mader KS, Schneider P, Müller R, Stampanoni M. A quantitative framework for the 3D characterization of the osteocyte lacunar system. Bone. 2013;57(1):142–54.
